# The role of the angular gyrus in semantic cognition – A synthesis of five functional neuroimaging studies

**DOI:** 10.1101/2021.12.21.473704

**Authors:** Philipp Kuhnke, Curtiss A. Chapman, Vincent K.M. Cheung, Sabrina Turker, Astrid Graessner, Sandra Martin, Kathleen A. Williams, Gesa Hartwigsen

## Abstract

Semantic knowledge is central to human cognition. The angular gyrus (AG) is widely considered a key brain region for semantic cognition. However, the role of the AG in semantic processing is controversial. Key controversies concern response polarity (activation vs. deactivation) and its relation to task difficulty, lateralization (left vs. right AG), and functional-anatomical subdivision (PGa vs. PGp subregions). Here, we combined the fMRI data of five studies on semantic processing (n = 172) and analyzed the response profiles from the same anatomical regions-of-interest for left and right PGa and PGp. We found that the AG was consistently deactivated during non-semantic conditions, whereas response polarity during semantic conditions was inconsistent. However, the AG consistently showed *relative* response differences between semantic and non-semantic conditions, and between different semantic conditions. A combined analysis across all studies revealed that AG responses could be best explained by separable effects of task difficulty and semantic processing demand. Task difficulty effects were stronger in PGa than PGp, regardless of hemisphere. Semantic effects were stronger in left than right AG, regardless of subregion. These results suggest that the AG is engaged in both domain-general task-difficulty-related processes and domain-specific semantic processes. In semantic processing, we propose that left AG acts as a “multimodal convergence zone” that binds different semantic features associated with the same concept, enabling efficient access to task-relevant features.

## Introduction

Semantic knowledge about objects, people and events in the world is crucial for core human cognitive abilities, such as object recognition and use, as well as language comprehension (Kiefer and Pulvermüller 2012; van Elk et al. 2014; Lambon Ralph 2014). The angular gyrus (AG) is widely considered a key brain region for semantic processing (for reviews, see Binder and Desai 2011; Seghier 2013; Price et al. 2015). This view is supported by meta-analyses of functional neuroimaging studies, which revealed consistent AG engagement for general semantic contrasts (e.g., words vs. pseudowords; Binder et al. 2009; Jackson 2021). Moreover, transcranial magnetic stimulation (TMS) studies indicate a causal role of left AG in general semantic processing (Sliwinska et al. 2015; Davey et al. 2015; Hartwigsen et al. 2016). It has been proposed that the AG acts as a cross-modal convergence zone that integrates semantic features related to various sensory-motor modalities (Binder 2016; Fernandino et al. 2016; Kuhnke et al. 2020b). This theory is corroborated by the AG’s proximity to and connectivity with several sensory-motor cortices (Bonner et al. 2013; Binder and Fernandino 2015; Kuhnke et al. 2021).

However, the role of the AG in semantic processing is controversial. Key controversies concern response polarity and its relation to task difficulty, lateralization, and functional-anatomical subdivision. First, the AG often shows deactivation (rather than positive activation) during both semantic and non-semantic tasks, as compared to a resting baseline (Lambon Ralph et al. 2016; Humphreys et al. 2021). This response profile is in line with the AG’s involvement in the default mode network (DMN), a set of brain regions that are functionally interconnected during rest and deactivated during attention-demanding tasks (Buckner et al. 2008; Raichle 2015). Notably, the amount of AG deactivation seems to be related to task difficulty (e.g., as measured by response times), where the AG shows less deactivation for easier conditions (Humphreys and Lambon Ralph 2017). This pattern has been observed for both semantic and non-semantic tasks, suggesting a domain-general, rather than semantics-specific role of the AG (Humphreys et al. 2015, 2021). Crucially, general semantic contrasts often compare easy vs. hard conditions (as in words vs. pseudowords). Therefore, it is unclear whether relative increases in AG activity for these contrasts indeed reflect semantic processing or merely domain-general task difficulty (Humphreys et al. 2021).

Second, it is unknown whether the left and right AG have the same or different functions in semantic cognition. In the meta-analysis of Binder et al. (2009), both left and right AG showed consistent engagement in semantic processing, albeit left AG showed a higher activation likelihood than right AG. In contrast, a more recent meta-analysis found exclusive recruitment of the left AG, but not right AG, across neuroimaging studies (Jackson 2021). Indeed, some individual studies reveal selective semantic effects in the left, but not right AG (Price et al. 2016; Kuhnke et al. 2020b), suggesting a potential functional lateralization. However, the response profiles of left and right AG in semantic and non-semantic tasks have not been systematically compared.

Finally, it remains unclear whether the AG constitutes a functional unit or comprises multiple functional subdivisions. The AG can be anatomically subdivided into two distinct cytoarchitectonic subregions: an anterior subregion *PGa*, and a posterior subregion *PGp* (Caspers et al. 2006, 2008). As function is generally assumed to follow cytoarchitecture (Fedorenko and Kanwisher 2009), it seems likely that PGa and PGp correspond to two functionally distinct areas. Functional subdivisions of the AG have been suggested previously (Seghier 2013). For example, a neuroimaging meta-analysis found consistent engagement of left dorsal AG for semantic tasks with an increased executive demand, whereas ventral AG was insensitive to control demands (Noonan et al. 2013). Therefore, the “controlled semantic cognition” account proposes that left dorsal AG supports the controlled retrieval of semantic representations, rather than semantic representation *per se* (Jefferies 2013). However, it is unclear to what extent these previous subdivisions correspond to the PGa vs. PGp distinction, and the response profiles of PGa and PGp have not been directly compared during semantic and non-semantic tasks.

To address these issues, we combined and re-analyzed the fMRI data of 5 different studies on semantic processing from our laboratory (Kuhnke et al. 2020b; Chapman and Hartwigsen 2021; Graessner et al. 2021; Turker et al. 2021; Martin et al. 2021). For each study, we investigated the response profiles of the same anatomical regions-of-interest (ROIs) for left and right PGa and PGp. Specifically, we asked whether each AG subregion shows significant activation or deactivation, as compared to the resting baseline, during semantic and non-semantic conditions. Moreover, we tested for relative activity differences between semantic and non-semantic conditions, as well as between different semantic conditions to probe the AG’s sensitivity to semantic variables. In a combined linear-mixed-model analysis across all studies, we then investigated whether the response profile of each AG subregion could be best explained by task difficulty, semantic processing demand (i.e., whether the task involves semantic processing), or both.

The view that the AG is a domain-general region showing task-difficulty-related deactivation (Lambon Ralph et al. 2016; Humphreys et al. 2021) would predict that the AG is consistently deactivated during both semantic and non-semantic conditions, and that the level of AG activity can be explained by task difficulty alone. In contrast, the view that the AG is engaged in semantic processing (Seghier 2013; Binder and Fernandino 2015; Kuhnke et al. 2020b) would predict that AG responses cannot be explained by task difficulty alone, but it is crucial to consider semantic processing demand. Regarding laterality, we hypothesized that left AG might show stronger semantic effects than right AG, given that left AG showed more consistent engagement in meta-analyses of semantic processing (Binder et al. 2009; Jackson 2021). Regarding functional subdivision of the AG, some previous work would predict that PGa is involved in domain-general task-difficulty-related processes, whereas PGp is engaged in semantic processing (Jefferies 2013; Noonan et al. 2013). Other research suggests that both PGa and PGp are involved in semantic processing, albeit with different semantic roles (Seghier 2013).

## Materials and Methods

### Studies

We combined the fMRI data of 5 different studies on semantic processing from our laboratory (Kuhnke et al. 2020b; Chapman and Hartwigsen 2021; Graessner et al. 2021; Turker et al. 2021; Martin et al. 2021). Table 1 presents an overview of all studies. Every study acquired blood oxygenation level dependent (BOLD) functional images in young and healthy human adults using a 3T MRI scanner. The total number of participants across all studies was 172. 36 participants participated in multiple studies, yielding 111 unique participants. This overlap was taken into account in combined analyses. Written informed consent was obtained from all participants. All studies were performed according to the guidelines of the Declaration of Helsinki and approved by the local ethics committee of the University of Leipzig. For further details on each study’s experimental design, measurement procedure and processing, please see the individual publications.

**Table 1.** Overview of all studies.

|  | Study | N | Age range | Tasks | Stimulus type | Stimulus modality | Stimulus conditions | Example trials |
| --- | --- | --- | --- | --- | --- | --- | --- | --- |
| **A** | **Chapman and Hartwigsen (2021)** | 41 | 21-35 years | taxonomic judgment; thematic judgment; scrambled judgment | pairs of line drawings (comprehension) | visual | **taxonomically related, thematically related & unrelated images;** *congruent & incongruent scrambled images* | taxonomic: moose & gorilla: taxonomically related?; thematic: monkey & banana: thematically related?; scrambled: does left image mirror right? |
| **B** | **Graessner et al. (2021)** | 33 | 19-34 years | explicit (meaningfulness judgment); implicit (lexical status comparison) | spoken 2-word phrases (comprehension) | auditory | **meaningful, anomalous, pseudoword phrases; single words** | explicit: "anxious horse" - meaningful?; implicit: "anxious gufel" - same lexical status? |
| **C** | **Kuhnke et al. (2020b)** | 40 | 19-33 years | lexical decision; sound judgment; action judgment | written words (comprehension) | visual | **words with a low or high association to sounds & actions;** *pseudowords* | lexical decision: "Telephone" - word?; sound judgment: "Telephone" - strongly associated with sounds?; action judgment: "Telephone" - strongly associated with actions? |
| **D** | **Martin et al. (2021)** | 30 | 21-34 years | overt semantic fluency; overt counting | written category names (production) | visual | **easy & difficult semantic categories;** *easy (forward) & difficult (backward) counting* | easy semantic categories: colors, body parts, animals; difficult semantic categories: flowers, metals, fish |
| **E** | **Turker et al. (2021)** | 28 | 18-29 years | overt reading | written words (production) | visual | simple (2-syllable) & complex (4-syllable) **words & pseudowords** | read aloud: "Firma", "Bürgermeister", "Molte", "Perautsarung" |

Semantic conditions are highlighted in bold font; non-semantic conditions are highlighted in italics.

For all subsequent analyses, each study’s fMRI data were re-preprocessed to match voxel size (2.5 mm^3^) and smoothing kernel size (5 mm^3^ FWHM) in Montreal Neurological Institute (MNI) space. Moreover, we ensured that no study removed response time (RT) related activity from their data as a central question of the present study is whether AG responses can be explained by task difficulty.

### Behavioral analyses

Behavioral analyses for each study focused on error rates and mean RTs for correct trials. Statistical inference was performed using repeated-measures ANOVAs, corrected for non-sphericity using the Huynh-Feldt method. Significant interactions were resolved using post-hoc paired-tests. P-values were corrected for multiple comparisons using the Bonferroni-Holm method.

### ROI analyses

For each study, we extracted the response profiles from the same anatomical regions-of-interest (ROIs) for left and right PGa and PGp. Anatomical ROIs were defined as the maximum probability maps for these cytoarchitectonic areas (Caspers et al. 2006, 2008) provided by the *SPM Anatomy* toolbox (Eickhoff et al. 2005, 2006). For each experimental condition and participant, we estimated percent signal change (PSC) as compared to the resting baseline using the *MarsBaR* toolbox (Brett et al. 2002). For statistical inference, we first performed one-sample t-tests on each experimental condition to test for significant activation or deactivation from the resting baseline. Second, to investigate relative activity differences between experimental conditions, we ran repeated-measures ANOVAs, correcting for non-sphericity using the Huynh-Feldt method. Significant interactions were resolved via step-down ANOVAs and post-hoc paired t-tests. P-values were corrected for multiple comparisons using the Bonferroni-Holm method.

### Linear-mixed-model analysis across all studies

To examine effects of semantic processing and task difficulty on AG activation, we employed a linear-mixed-effects modeling approach across all studies. First, we used a goodness-of-fit comparison to determine the best-fitting model using the Akaike Information Criterion (AIC), where a model was considered meaningfully more informative than others if it decreased the AIC by at least two points (Burnham and Anderson 2004). To reduce overfitting, AIC takes into account model complexity by penalizing models with more parameters. Models predicted PSC across ROIs using various fixed and random effects. The optimal model for our data in terms of AIC was determined in a stepwise fashion, first determining the optimal random effects structure (i.e., individual random effects of task and condition, nested effects, etc.); next determining which measure(s) of task difficulty (response times or error rates) better predicted PSC; and finally determining which interactions of task difficulty, semantic processing demand, and ROI were optimal. In describing model selection, we name the best model and then describe the alternative models and their difference in AIC (ΔAIC).

Models were run using the *lme4* package (Bates et al. 2015) in *R* (version 4.1.1; R Core Team), and model AIC comparisons were performed using the *bbmle* package (Bolker 2020). Significance of effects from the optimal model and further models was derived using the Satterthwaite approximation for degrees of freedom from the *R* package *lmerTest* (Kuznetsova et al. 2017), and predicted plots of interactions were created using the *R* packages *sjPlot* (Lüdecke 2021) and *ggeffects* (Lüdecke 2018). Significant interactions from the optimal model were further investigated by specifying contrasts with the *R* package *hypr* (Rabe et al. 2020) in the manner suggested by Schad et al. (2020). In addition, versions of the optimal model were also run in each ROI individually to test effects of task difficulty and semantics in each ROI without assuming homogeneity of variance. These models were identical to the optimal model but excluded the ROI term.

### Results

#### Behavioral and ROI analyses for each study

##### Study A: Chapman and Hartwigsen (2021)

In the study by Chapman and Hartwigsen (2021), 41 young and healthy adults had to decide whether pairs of object line drawings were taxonomically related (taxonomic task), or thematically related (thematic task). In both tasks, the drawings could be taxonomically related (e.g., *monkey* and *moose*), thematically related (e.g., *monkey* and *banana*), or unrelated (e.g., *monkey* and *telephone*). In a non-semantic control task, participants decided whether pairs of scrambled versions of images from the semantic tasks mirrored each other.

Behavioral analyses revealed main effects of TASK and CONDITION, as well as TASK x CONDITION interactions for both error rates and mean RTs (Figure 1A; Tables S1-S2 for statistics). Error rates were higher for thematic than taxonomic judgments, but did not differ between semantic and non-semantic tasks. In the taxonomic task, but not in the thematic task, error rates were higher for taxonomic than thematic and unrelated conditions. Mean RTs were longer in the non-semantic than semantic tasks, and in the thematic than taxonomic task. In the taxonomic, but not in the thematic task, taxonomic and thematic conditions produced longer RTs than unrelated conditions.

**Figure 1.**
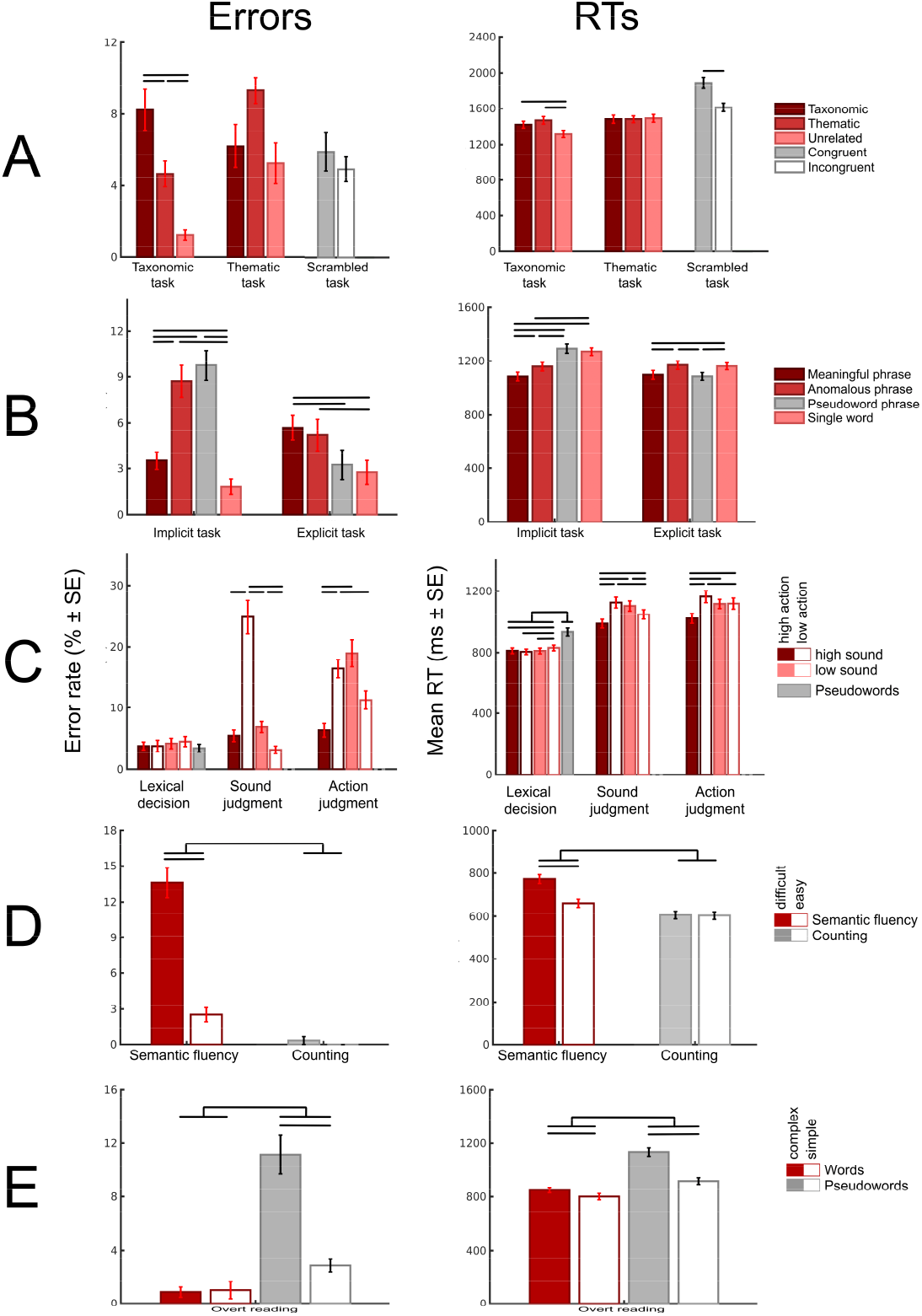
Behavioral results for all studies. Error rates and mean RTs for correct trials are plotted for each experimental condition (grouped by task). Error bars represent standard error of the mean. Semantic conditions are highlighted in red; non-semantic conditions are in gray. Black bars illustrate significant differences between conditions (p < 0.05 Bonferroni-Holm corrected).

All AG-ROIs were deactivated in the non-semantic scrambled task and exhibited higher activity for semantic than non-semantic tasks (Figure 2A; Tables S11-S13 for statistics). However, the four AG-ROIs showed distinct response profiles during the semantic tasks (ROI x TASK x CONDITION interaction). Left PGp was the only AG-ROI to show positive activation, which occurred for thematic pairs during thematic judgments. Both left AG subregions showed relatively higher activity for thematic pairs than taxonomic and unrelated pairs across semantic tasks. Right PGa was deactivated for all conditions, with less deactivation for taxonomic and thematic than unrelated pairs across tasks. Finally, right PGp did not show activation differences from resting baseline in the semantic tasks. However, right PGp was relatively more engaged for thematic than taxonomic and unrelated pairs across tasks.

**Figure 2.**
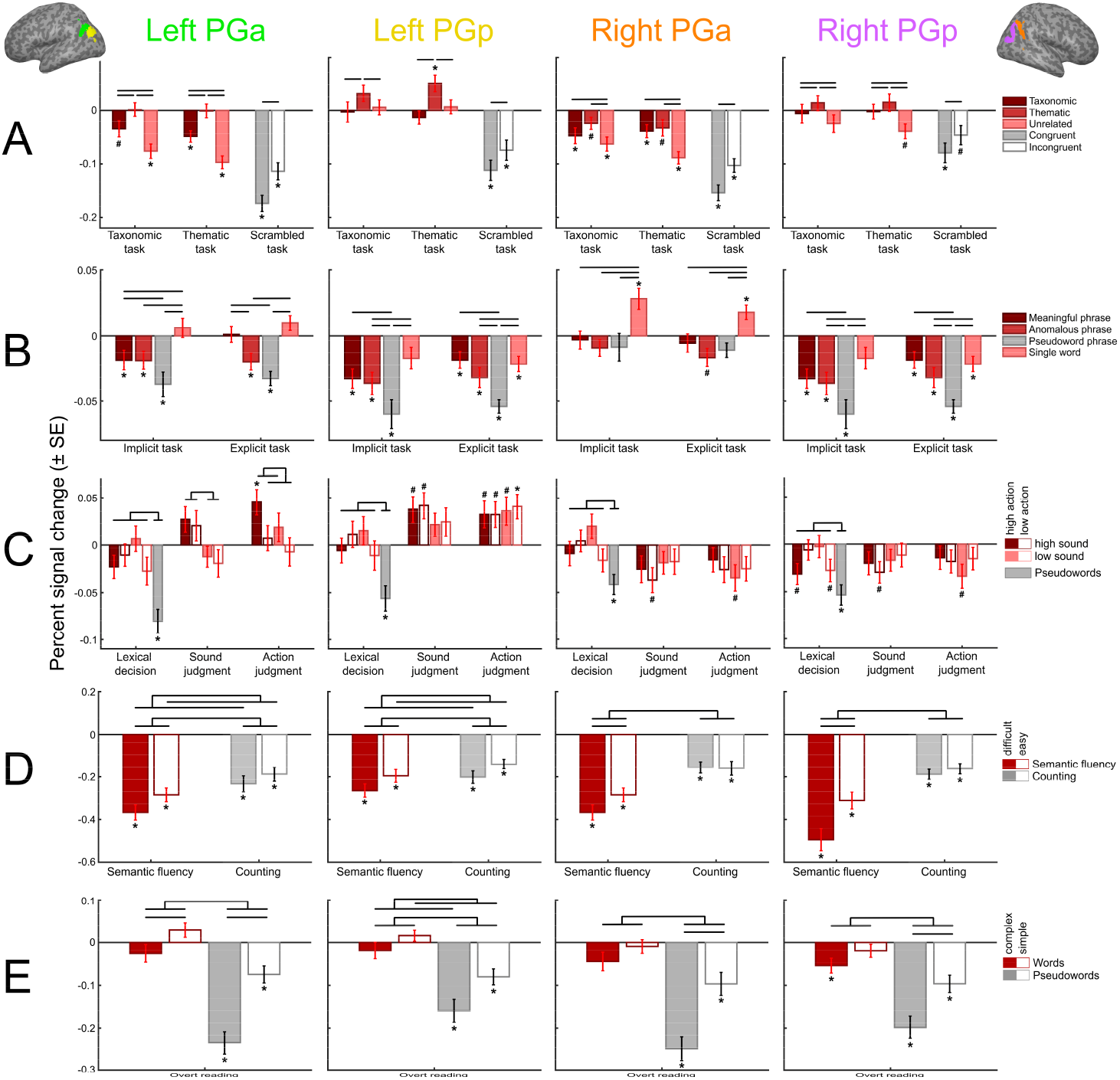
Response profiles of each AG subregion for every study. Mean percent signal change as compared to the resting baseline is plotted for each experimental condition (grouped by task). Error bars represent standard error of the mean. Semantic conditions are highlighted in red; non-semantic conditions are in gray. Black bars illustrate significant differences between experimental conditions. *: p < 0.05 (Bonferroni-Holm corrected); #: p < 0.05 (uncorrected).

##### Study B: Graessner et al. (2021)

In Graessner et al. (2021), 33 participants performed explicit (meaningfulness judgment) and implicit (lexical status judgment) semantic tasks on auditorily presented meaningful phrases (e.g., “fresh apple”), anomalous phrases (“awake apple”) and pseudoword phrases (“fresh gufel”), as well as single words (“apple”).

Behavioral analyses yielded main effects of TASK and CONDITION, and TASK x CONDITION interactions for both error rates and mean RTs (Figure 1B; Tables S3-S4). Error rates were higher in the implicit than explicit task. In the implicit task, error rates were higher for anomalous and pseudoword phrases than meaningful phrases and single words. In the explicit task, error rates were higher for meaningful and anomalous phrases than pseudoword phrases and single words. RTs were slower for anomalous than meaningful phrases in both tasks. In the implicit task, pseudoword phrases and single words yielded longer RTs than anomalous phrases and meaningful phrases. In the explicit task, anomalous phrases and single words led to longer RTs than meaningful and pseudoword phrases.

Left PGa showed a distinct response profile from the other AG-ROIs (ROI x TASK x CONDITION interaction) (Figure 2B; Tables S14-S16). In both tasks, left PGa was significantly deactivated for anomalous and pseudoword phrases. Pseudoword phrases elicited stronger deactivation than meaningful and anomalous phrases. However, selectively in the explicit task (i.e., not in the implicit task) and selectively in left PGa, meaningful phrases induced relatively higher activity than anomalous phrases (TASK x CONDITION interaction).

All other AG-ROIs showed a main effect of CONDITION. Left and right PGp were significantly deactivated for all phrases in both tasks. Pseudoword phrases induced stronger deactivation as compared to meaningful phrases in both regions, and as compared to anomalous phrases in left but not right PGp.

Right PGa was significantly activated for single words, but not for phrases, in both tasks. Single words produced higher activity than all phrase types (which did not significantly differ).

##### Study C: Kuhnke et al. (2020b)

In the study by Kuhnke et al. (2020b), 40 participants performed three different tasks—lexical decision, sound judgment, and action judgment—on written words with a high or low association to sounds and actions (e.g., “guitar” was a high sound–high action word). Pseudowords acted as a non-semantic control condition.

Behavioral analyses showed main effects of TASK, SOUND, ACTION, as well as TASK x SOUND x ACTION interactions for both error rates and mean RTs (Figure 1C; Tables S5-S6). Error rates and RTs were lower in the lexical decision than semantic judgment tasks. In the lexical decision task, error rates did not significantly differ between conditions. Pseudowords yielded higher RTs than words, and low sound–low action words produced higher RTs than the other word types. In the sound judgment task, high sound–low action words produced higher error rates than all other word types. High sound–low action and low sound–high action words were associated with higher RTs than the other words. In the action judgment task, high sound–low action and low sound–high action words yielded higher error rates than the other word types. High sound–low action words produced slower RTs than the other words.

In the lexical decision task, none of the AG-ROIs showed significant (de-)activation for the four word types, but all regions demonstrated strong deactivation for pseudowords that differed significantly from activity for words (Figure 2C; Tables S17-S19). In the semantic judgment tasks, the four AG-ROIs exhibited distinct response profiles (ROI x TASK x SOUND and ROI x TASK x ACTION interactions). Left PGa showed relatively higher activity for high-than low-sound words during sound judgments. During action judgments, left PGa was more strongly engaged for high-than low-action words. High-sound high-action words elicited significant positive activation as compared to the resting baseline.

In contrast, left PGp showed a main effect of TASK, driven by higher positive activation for all word types during semantic judgments than lexical decisions. (Sound and action judgments did not significantly differ.)

Contrary to the left AG regions, right PGa and PGp showed no positive activation vs. rest in any condition, and no significant activity differences between semantic conditions.

##### Study D: Martin et al. (2021)

In the study by Martin et al. (2021), 30 healthy adults performed an overt semantic fluency task on easy categories (e.g., colors, animals) and difficult categories (e.g., metals, flowers), as well as an overt counting task with easy (forward) and difficult (backward) conditions.

Behavioral analyses revealed main effects of TASK and DIFFICULTY, as well as TASK x DIFFICULTY interactions for both error rates and mean RTs (Figure 1D; Tables S7-S8). The semantic fluency task yielded higher error rates and RTs than the counting task. Selectively in the semantic fluency task, but not in the counting task, difficult conditions produced significantly higher error rates and RTs than easy conditions.

All AG-ROIs were significantly deactivated during all conditions (Figure 2D; Tables S20-S22). Moreover, all AG regions exhibited stronger deactivation during the semantic fluency than counting task (main effect of TASK).

However, left and right AG regions showed a striking difference in their relative responses (ROI x TASK x DIFFICULTY interaction). Left PGa and PGp were more deactivated for difficult than easy conditions across both tasks (main effect of DIFFICULTY). In contrast, right PGa and PGp showed a selective difficulty effect in the semantic fluency task, but not in the counting task (TASK x DIFFICULTY interaction).

##### Study E: Turker et al. (2021)

In Turker et al.’s (2021) study, 28 participants read aloud simple (two-syllable) and complex (four-syllable) words (e.g., “Firma”, “Bürgermeister”) and pseudowords (e.g., “Molte”, Perautsarung”).

Behavioral analyses revealed main effects of WORD and COMPLEXITY, as well as WORD x COMPLEXITY interactions for both error rates and mean RTs (Figure 1E; Tables S9-S10). Both error rates and mean RTs were higher for pseudowords than words. Selectively for pseudowords, but not for words, complex stimuli yielded higher error rates than simple stimuli. Complex words and pseudowords both yielded higher RTs than simple stimuli, but this complexity effect was larger for pseudowords than words.

All AG-ROIs were significantly deactivated for both simple and complex pseudowords (Figure 2E; Tables S23-S25). Words did not induce significant activation or deactivation, as compared to the resting baseline. As an exception, right PGp was deactivated for complex words.

However, left and right AG-ROIs exhibited distinct relative responses (ROI x WORD x COMPLEXITY interaction). Left PGa and PGp showed a complexity effect (complex vs. simple) for both words and pseudowords. In contrast, right PGa and PGp showed a selective complexity effect for pseudowords, but not for words.

### Commonalities and differences between studies

In all studies, all AG-ROIs were consistently deactivated during non-semantic conditions. As the only exception, right PGa did not show a significant difference from resting baseline in study B (Graessner et al. 2021).

In contrast, response polarity during semantic conditions was inconsistent between ROIs, studies, and conditions. The same AG subregion could show both positive activation and deactivation in different studies, or even in different conditions of the same study. Notably, both study A (Chapman and Hartwigsen 2021) and study C (Kuhnke et al. 2020b) found significant positive activation in left AG during explicit semantic judgment tasks.

Relative activity differences between conditions were more consistent. In all studies, all AG-ROIs showed significant activity differences between semantic and non-semantic conditions.

This semantic effect was generally driven by relatively higher activity for semantic than non-semantic conditions. The only exception was study D (Martin et al. 2021), where a semantic fluency task induced relatively lower activity than a counting task. It is questionable, however, whether counting is indeed a non-semantic task (see Discussion).

Crucially, left AG not only showed relative activity differences between semantic and non-semantic conditions, but also between different semantic conditions. The same pattern was sometimes, but not always, observed in right AG (e.g., non-significant differences in studies C and E).

Finally, in addition to functional differences between left and right AG, we found distinct response profiles for the cytoarchitectonic subregions PGa and PGp. Out of all AG-ROIs, left PGa seemed to exhibit the strongest sensitivity to fine-grained semantic manipulations. For example, in study B (Graessner et al. 2021), left PGa was the only AG-ROI that showed relatively higher activity for meaningful phrases (e.g., “fresh apple”) than anomalous phrases (e.g., “awake apple”) during explicit meaningfulness judgments. In study C (Kuhnke et al. 2020b), left PGa was the only AG subregion that showed a selective response to task-relevant semantic features, that is, higher activation for high-than low-sound words during sound judgments and for high-than low-action words during action judgments.

### Linear-mixed-model analysis across all studies

To investigate whether the response patterns of each AG subregion could be best explained by task difficulty, semantic processing demand (i.e., whether the task involves semantic processing), or both, we performed a linear-mixed-model analysis across all studies.

First, to determine the optimal random effects structure for our model, we tested a baseline null model predicting percent signal change (PSC) with only the mean signal and random intercepts for participants. The models being compared always included random intercepts for participants, but they differed in the task and condition random effects. Results showed that the optimal random effects structure included a random intercept for participant and a random intercept for condition nested within task (Table 2A).

**Table 2.**
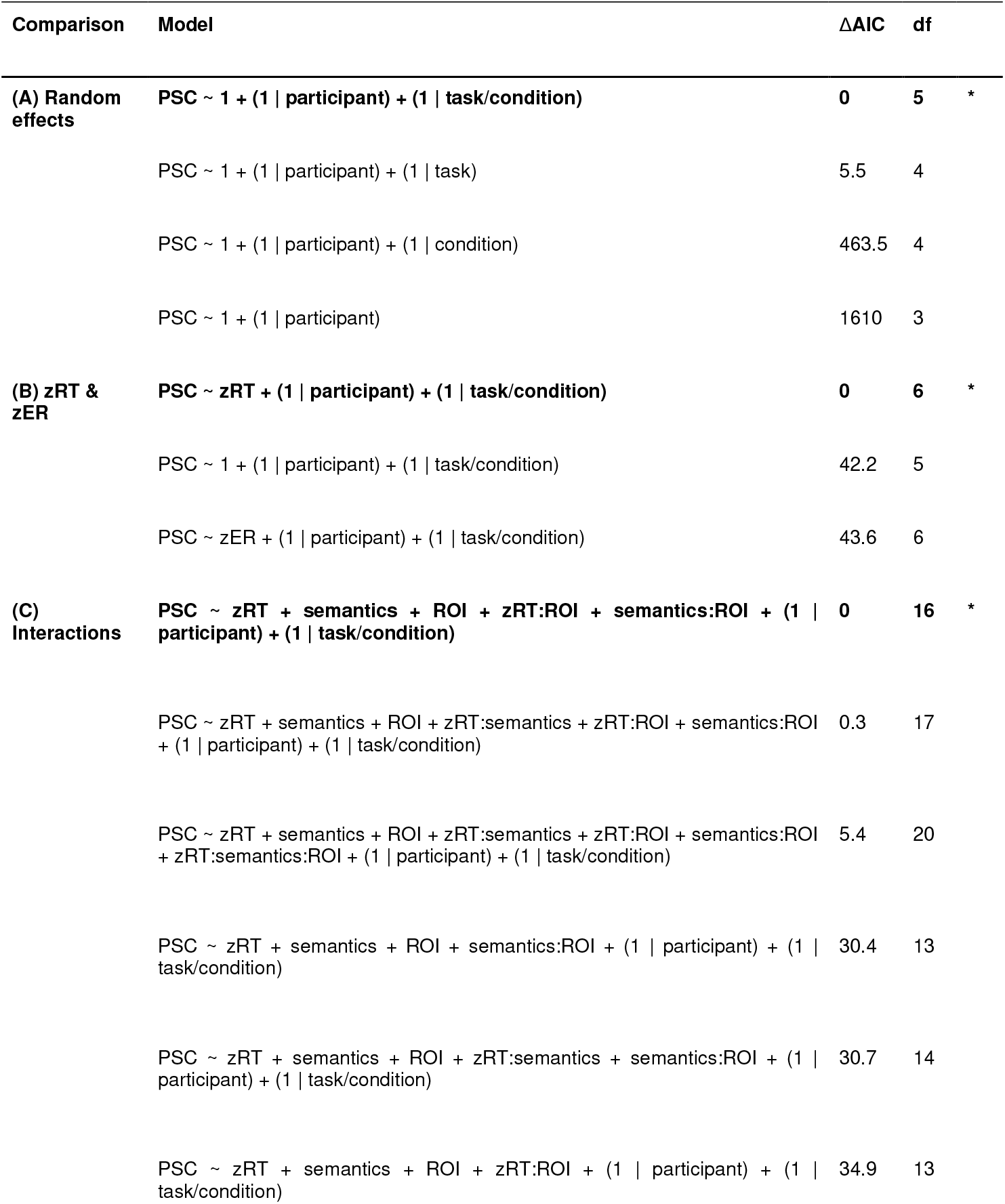
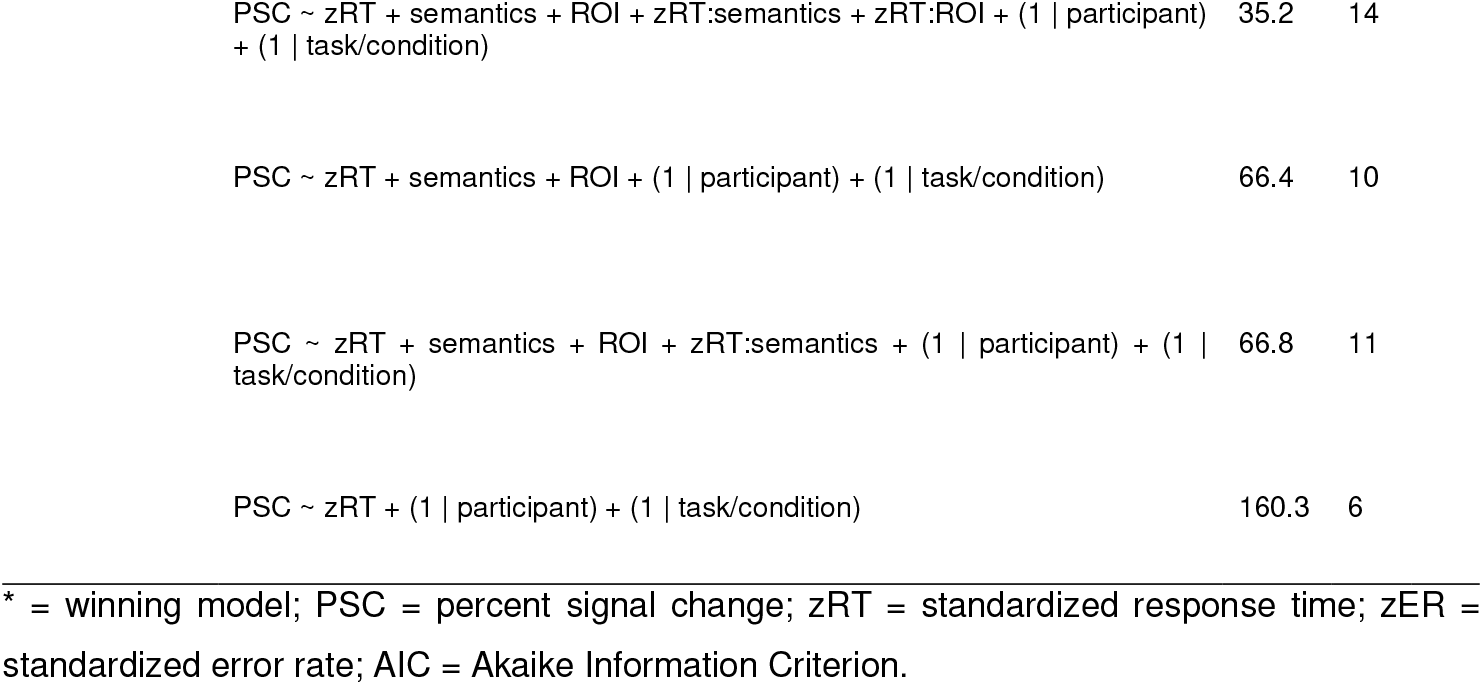
AIC comparisons to determine the optimal model predicting AG activity.

Next, we determined whether standardized error rates (zER) and/or standardized response times (zRT) were good predictors of PSC. Models with a single fixed effect—zER or zRT—were compared to our null intercept model. Results showed that including zRT as a fixed effect improved model fit, but the model including zER was not better than the null model (Table 2B). Thus, zRT was included as a fixed effect in further models and zER was excluded.

Finally, we compared models with three fixed effects—zRT, semantics, and ROI—and all combinations of interactions between the three. The model with the highest explanatory value included a 2-way interaction of zRT x ROI as well as a 2-way interaction of semantics x ROI (Table 2C). The ANOVA table for this optimal model is shown in Table 3.

**Table 3.** ANOVA table for the optimal model of AG responses across studies.

| Factor | SS | MSE | DF | Den. DF | F | p |
| --- | --- | --- | --- | --- | --- | --- |
| zRT | 0.37 | 0.37 | 1 | 4822.9 | 45.68 | < 0.001 |
| Sem | 0.09 | 0.09 | 1 | 27.7 | 11.50 | 0.002 |
| ROI | 0.35 | 0.12 | 3 | 5187.4 | 14.67 | < 0.001 |
| zRT x ROI | 0.29 | 0.10 | 3 | 5187.4 | 12.18 | < 0.001 |
| Sem x ROI | 0.33 | 0.11 | 3 | 5187.4 | 13.67 | < 0.001 |

SS = sum of squares; MSE = mean squared error; DF = degrees of freedom; Den = denominator; Sem = effect of semantics.

Although the next best-fitting model was within 2 ΔAIC from the optimal model, suggesting no meaningful difference, the additional zRT x Semantics variable implies that this model may be overfit. Moreover, an ANOVA on this more complex model revealed no significant effect of the zRT x Semantics interaction, whereas the zRT x ROI and semantics x ROI interactions remained significant (Table S26). These results indicate that the zRT x Semantics interaction was redundant to explain AG responses. In addition, two separate linear-mixed-model analyses on semantic and non-semantic conditions also revealed main effects of zRT, ROI and zRT x ROI (Tables S32-S35), indicating that effects of zRT on AG activity were domain-general and independent of semantics. Therefore, we interpret results from the parsimonious optimal model.

A plot of the predicted interaction between zRT and ROI based on the optimal model (Figure 3A) suggested that the effect of zRT was strongest in PGa and weaker in PGp. A plot of the predicted interaction between semantics and ROI (Figure 3B) suggested that the effect of semantics was stronger in left than right AG-ROIs. These interactions were investigated further by adding specific ROI contrasts to the model (Table S27).

**Figure 3.**
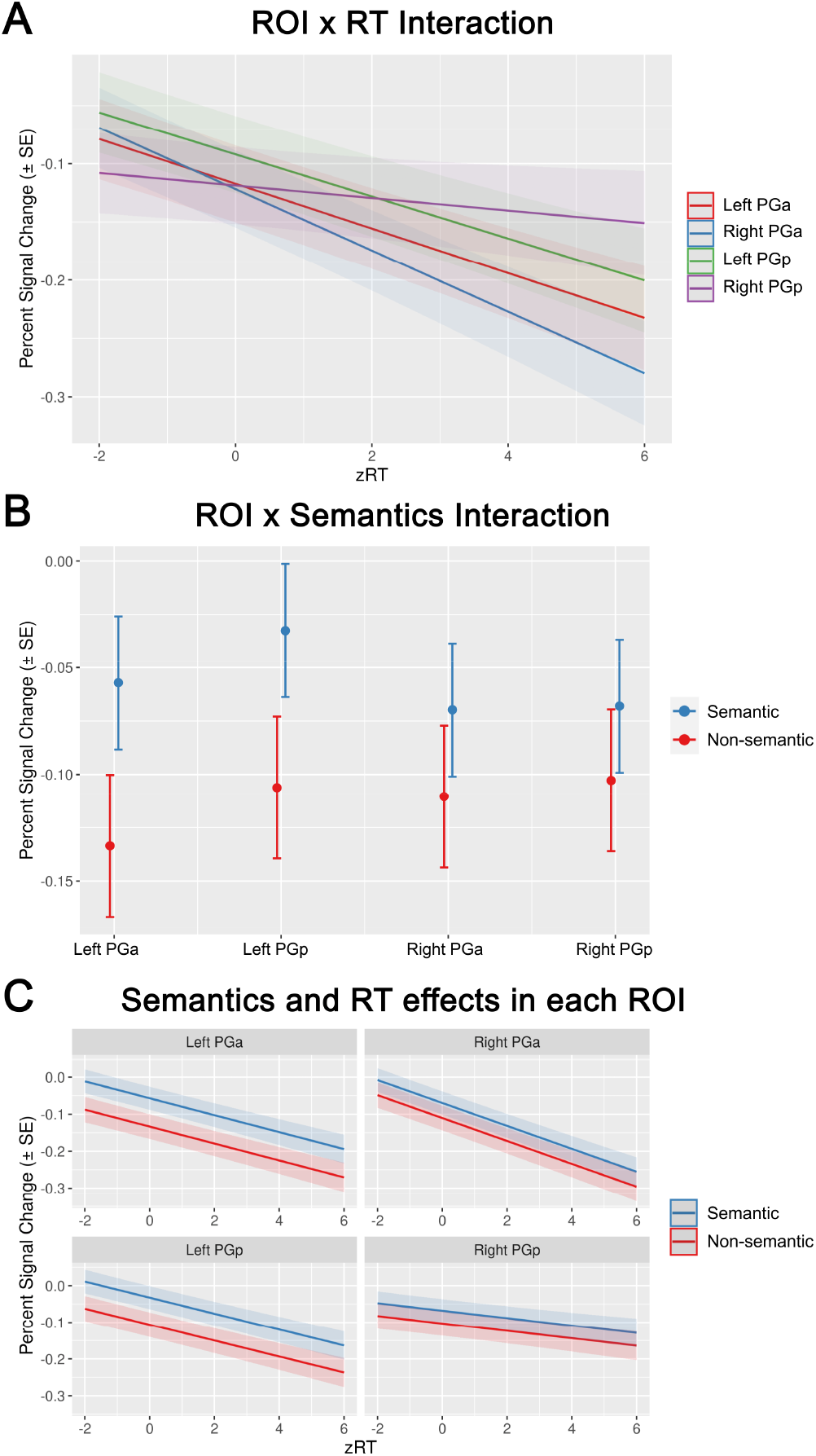
Results of the linear-mixed-model analysis. (A) Interaction of ROI and zRT on predicted percent signal change. (B) Interaction of ROI and semantics on predicted percent signal change. (C) Combined effects of semantics and zRT on predicted percent signal change in each ROI. Error bars and shaded areas represent standard error of the mean.

Specifically, the contrast model included contrasts of PGa vs. PGp and left vs. right ROIs. We found that zRT interacted with the PGa vs. PGp contrast (coeff = −0.022, p < 0.001) but not with the left vs. right contrast (coeff = −0.004, p = 0.47), providing evidence that the zRT x ROI contrast was due to a larger effect of zRT in PGa than PGp, regardless of hemisphere. Moreover, effects of semantics interacted with the left vs. right contrast (coeff = 0.075, p < 0.001), but not with the PGa vs. PGp contrast (coeff = 0.009, p = 0.46). This indicates that the semantics x ROI interaction was driven by a stronger effect of semantics in left than right AG, regardless of subregion.

Finally, we examined the zRT and semantic effects in each ROI individually (Figure 3C; Tables S28-S31). This step-down analysis allowed us to investigate the effects of task difficulty and semantics without assuming homogeneity of variance in each ROI. These models revealed that in addition to the independent effects of difficulty and semantics across all AG ROIs, each individual ROI showed independent main effects of difficulty (zRT) (Left PGa coefficient = −0.013, *p* = 0.015; Left PGp coefficient = −0.024, *p* < 0.001; Right PGa coefficient = −0.027, *p* < 0.001; Right PGp coefficient = −0.021, *p* < 0.001) and semantics (Left PGa coefficient = 0.073, *p* = 0.001; Left PGp coefficient = 0.057, *p* < 0.001; Right PGa coefficient = 0.051, *p* = 0.012; Right PGp coefficient = 0.044, *p* = 0.021).

Overall, the mixed-model analysis across all studies revealed that AG responses are best explained by separable effects of task difficulty (as measured by zRT) and semantic processing demand. AG activity decreased with increasing task difficulty. This task performance effect was larger in PGa than PGp, regardless of hemisphere. Moreover, AG activity was relatively higher for semantic than non-semantic conditions. This semantic effect was larger in left than right AG, regardless of subregion. The model intercept is negative, indicating that the AG is deactivated on average. However, AG activity was sometimes positive, especially during semantic conditions (see Figure S1).

## Discussion

This study tested the functional involvement of the angular gyrus (AG) in semantic cognition, focusing on three key issues: (1) response polarity (activation vs. deactivation) and its relation to task difficulty, (2) lateralization (left vs. right AG), and (3) functional-anatomical subdivision (PGa vs. PGp). To this end, we combined and re-analyzed the fMRI data of five studies on semantic processing from our laboratory. For each study, we extracted the response profiles from the same anatomical regions-of-interest (ROIs) for left and right PGa and PGp.

We found that the AG was consistently deactivated during non-semantic conditions, as compared to the resting baseline. In contrast, response polarity was inconsistent during semantic conditions, involving both deactivation and activation in different studies, conditions and AG-ROIs. However, we consistently found *relative* response differences between semantic and non-semantic conditions, as well as between different semantic conditions. These effects were always observed in left AG, and often but not always in right AG.

A combined linear-mixed-model analysis across all studies revealed that these response profiles could be best explained by both task difficulty (as measured by response times; RT) and semantic processing demand (Figure 4). AG activity decreased with increasing task difficulty and was relatively higher for semantic than non-semantic conditions. Difficulty effects were stronger in PGa than PGp, irrespective of hemisphere. Semantic effects were stronger in left than right AG, regardless of subregion.

**Figure 4.**
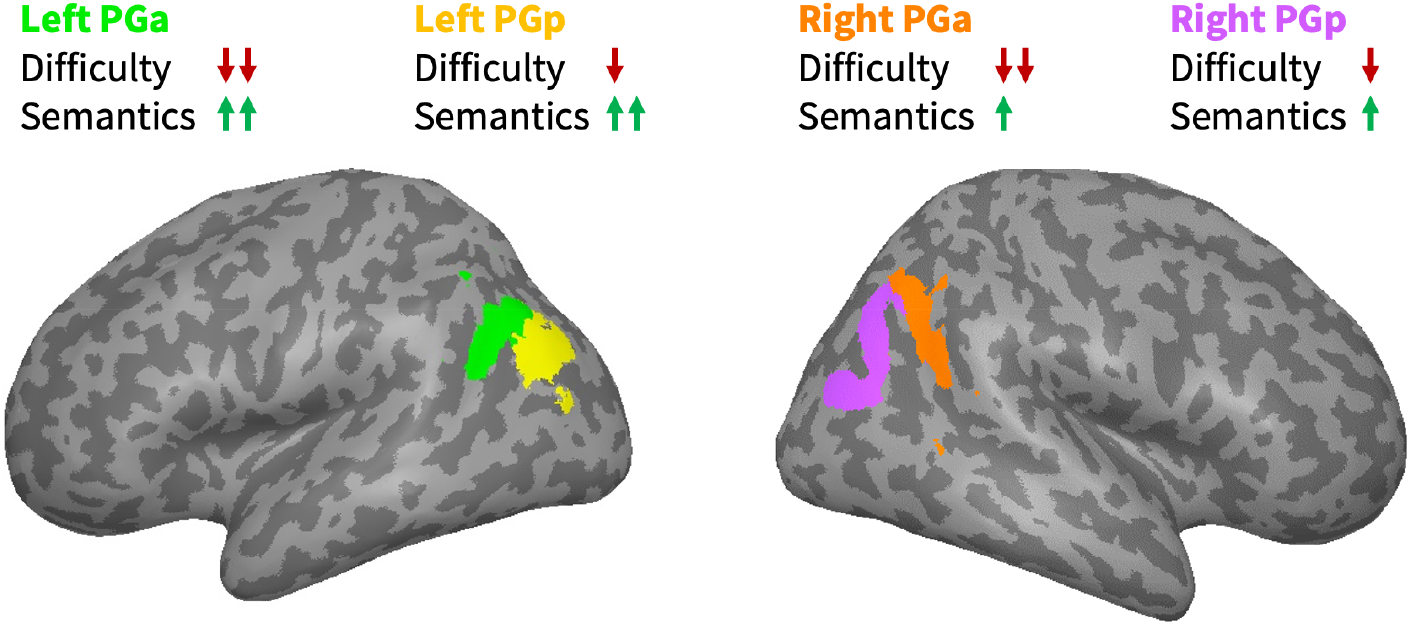
Overview of key findings. In all AG regions, neural activity depends on both task difficulty and semantic processing demand. Activity decreases with increasing difficulty, and is relatively higher for semantic than non-semantic conditions. Difficulty effects are stronger in PGa than PGp, regardless of hemisphere. Semantic effects are stronger in left than right AG, regardless of subregion.

### Theories of AG Function

Our findings support the view that the AG is engaged in semantic processing (Binder and Desai 2011; Seghier 2013; Kuhnke et al. 2020b), while they oppose the view that the AG is exclusively a domain-general region showing task-difficulty-related deactivation (Lambon Ralph et al. 2016; Humphreys et al. 2021). The domain-general view would have predicted that the AG is consistently deactivated during both semantic and non-semantic conditions, and any response differences between semantic and non-semantic conditions can be completely explained by task difficulty differences. In contrast, the semantics view would have predicted that AG responses cannot be explained by task difficulty alone, but it is crucial to consider semantic processing demand (i.e., whether the task involves semantic processing).

In line with the domain-general view, we found consistent deactivation of the AG during non-semantic conditions. Moreover, AG activity was related to task difficulty in both semantic and non-semantic conditions. However, response polarity during semantic conditions was inconsistent and involved *positive* activations (e.g., in studies A and C). Crucially, contrary to the domain-general view and in support of the semantics view, AG activity levels could not be explained task difficulty alone, but semantic processing demand proved essential: Models based on task difficulty alone were substantially outperformed by models that also included semantic processing demand. The optimal model included two-way interactions of ROI x RT and ROI x Semantics, but no RT x Semantics interaction, indicating that effects of RT and semantics on activity levels in each AG-ROI were independent. These results strongly support the view that the AG is engaged in semantic processing.

### Response Polarity and Semantic Processing in the AG

The inconsistency of AG response polarity during semantic conditions suggests that responses vs. rest are unreliable evidence to assess the AG’s role in semantic processing, contrary to the arguments in some previous work (Humphreys et al. 2015, 2021). More generally, comparisons against rest are problematic as the resting baseline itself is not process-neutral (Stark and Squire 2001; Morcom and Fletcher 2007). “Resting” can involve mind wandering, autobiographical memory, as well as self-referential and introspective processes (Andrews-Hanna 2012). It is particularly problematic that all these processes may involve the retrieval of semantic information (Binder et al. 1999, 2009). Thus, the AG might be “deactivated” during attention-demanding tasks as the semantic processing that occurs during rest is interrupted (Seghier 2013). In other words, AG deactivation may indeed reflect its involvement in semantic processing. Under this view, it is not surprising that AG activity in our semantic conditions often did not significantly differ from the resting baseline. A semantic region would only be predicted to show positive activation (above rest) when the task involves semantic processing to a *greater* extent than during the resting state. Indeed, we found positive activation for some semantic conditions (e.g., in studies A and C).

Compared to absolute responses vs. rest, *relative* responses between different experimental conditions were much more consistent: The AG consistently showed differential activity between semantic and non-semantic conditions, and between different semantic conditions. This strongly suggests that the AG is sensitive to semantics. Moreover, these findings corroborate the view that relative responses between conditions, not absolute responses vs. rest, should constitute the main focus of neuroimaging studies on AG function (Stark and Squire 2001; Finn 2021).

The AG generally showed relatively *higher* activity for semantic than non-semantic conditions. The only exception was study D (Martin et al. 2021), where a semantic fluency task induced relatively lower activity than a counting task. Several possibilities may explain this result: First, counting involves the production of number words, which are often considered a type of abstract concept (Hauk and Tschentscher 2013; Desai et al. 2018). Thus, counting may indeed involve abstract semantic processes to a greater extent than a semantic fluency task on everyday object concepts (e.g., flowers, animals). This view is supported by the fact that counting is frequently employed in clinical contexts for pre-operative language mapping (Duffau et al. 1999, 2004). Second, the AG response pattern in study D may be mainly driven by task difficulty: Behavioral analyses revealed that the counting task was easier than the semantic fluency task. Therefore, the domain-general task difficulty effect may have overshadowed the semantic effect in this study.

Overall, our results suggest that AG responses are modulated by both stimulus characteristics and task demands. Regarding stimulus characteristics, the AG is more engaged for meaningful than meaningless stimuli, even when presented in the same task (e.g., words > pseudowords in studies B, C and E; related > unrelated object pairs in study A; meaningful > meaningless phrases in study B). Other stimulus variables known to modulate AG responses, such as concreteness (Binder et al. 2005), frequency (Graves et al. 2010) or familiarity (Woodard et al. 2007) were well-matched between conditions within each study, and potential between-study differences were controlled for in the random effects structure of our linear-mixed-effects model. Regarding task demands, the AG seems to show stronger activity in tasks that explicitly require the retrieval of semantic information (e.g., semantic judgments) than tasks that only implicitly probe semantic processing (e.g., lexical decision). Notably, a supplementary linear-mixed-model analysis that distinguished explicit and implicit semantic tasks yielded no improvement in model fit (Table S38), suggesting that the AG is equally involved in both types of task. However, fine-grained differences between explicit and implicit semantic tasks were observed in individual studies: In study B (Graessner et al. 2021), left PGa selectively showed higher activity for meaningful phrases (e.g., “fresh apple”) than anomalous phrases (e.g., “awake apple”) during explicit meaningfulness judgments, but not during implicit lexical comparisons. In study C (Kuhnke et al. 2020b), left PGa was selectively engaged for sound features of word meaning during sound judgments, and for action features during action judgments, but for neither during lexical decisions. Moreover, we observed positive activation (above rest) in the left AG in studies A and C when the task explicitly required the retrieval of individual semantic features. Taken together, these results suggest that left AG responds most strongly to task-relevant semantic information. This view is in line with theories that assume semantic processing to rely on a flexible, task-dependent architecture (Hoenig et al. 2008; Kemmerer 2015; Kuhnke et al. 2020b, 2021).

### Task Difficulty Effects in the AG

Several recent studies have claimed that semantic effects in the AG are likely to be an artifact of difficulty rather than semantic processing *per se* (Humphreys et al. 2015, 2021). That is, the authors point out that some typical semantic contrasts that reveal larger relative activation in the AG may be confounded with difficulty (e.g., words vs. pseudowords, concrete vs. abstract words). Indeed, multiple studies found that harder conditions yield stronger AG deactivation (Hahn et al. 2007; Humphreys et al. 2015). The current study reveals that task difficulty and semantic processing demand show *separable* effects in the bilateral AG: AG activity decreased with increasing task difficulty across semantic and non-semantic conditions. Independently, AG activity was relatively higher during semantic than non-semantic conditions. Notably, difficulty and semantic effects were also orthogonal in regional preference: Difficulty effects were stronger in PGa than PGp, regardless of hemisphere. Semantic effects were stronger in left than right AG, regardless of subregion. Thus, our results are consistent with the claim that the AG indexes task difficulty, but they provide strong evidence against the claim that semantic effects in the AG are explained by difficulty.

The separability of difficulty and semantic effects provides evidence that the AG is likely to have a domain-general role in addition to being sensitive to semantic processing. This view is supported by a previous fMRI study which revealed that the same regions of the default mode network (DMN), including AG, can show domain-general task difficulty effects (activation for pseudowords vs. words) and domain-specific semantic effects (decoding of high vs. low imageability words) (Mattheiss et al. 2018). Moreover, this view is in line with modern networkbased views of cognitive neuroscience, suggesting that the function of a brain region depends on its interactions with other areas in a given task (Seghier 2013). Under this view, the same region can have multiple different functions by virtue of being connected to different regions in different tasks (Bassett and Sporns 2017). Indeed, the AG exhibits particularly flexible functional connectivity in different cognitive tasks (Chai et al. 2016; Kuhnke et al. 2021). The AG seems to be involved in *both* domain-general task-difficulty-related processes and domain-specific semantic processes. These are not mutually exclusive.

Relatedly, it is important to consider whether semantics and task difficulty were correlated in our study, and if so, how the linear-mixed-model analysis could distinguish their effects on AG activity. Firstly, non-semantic conditions were not always harder than semantic conditions. For example, in study C (Kuhnke et al. 2020b), lexical decisions on pseudowords were easier than semantic judgments. Nonetheless, pseudowords induced the strongest AG deactivation. Secondly, the semantics and RT variables were not strongly correlated (Table S37), allowing for both variables to explain unique variance in AG responses. Finally, if semantic effects in the AG could be completely explained by RT, then a model based only on RT should be optimal. However, this was clearly not the case: The RT-only model was substantially outperformed by models that also included semantics. Overall, our results indicate that task difficulty and semantics explain separable parts of the variance in AG activity.

Notably, while the linear-mixed-model analysis across all studies indicated that effects of task difficulty and semantics on AG responses were independent, we found an interaction between semantics and difficulty in study D (Martin et al. 2021). Specifically, right AG showed a selective difficulty effect in the semantic fluency task, but not in the counting task. However, behavioral analyses also revealed a selective difficulty effect in the semantic fluency task, whereas forward (“easy”) and backward (“difficult”) counting did not differ in behavioral performance. Therefore, the AG response pattern corresponded to the behavioral pattern, consistent with the view that difficulty effects in the AG are domain-general.

### Lateralization

Left AG showed stronger semantic effects than right AG. Our combined analysis across all studies revealed that left AG exhibits larger activity differences between semantic and non-semantic conditions. In individual studies, left AG always showed activity differences between semantic and non-semantic conditions, as well as between different semantic conditions. Right AG also often showed these effects, but not always. For example, in study C (Kuhnke et al. 2020b), left but not right AG was engaged for sound and action features of word meaning. In study E (Turker et al. 2021), only left but not right AG was sensitive to word complexity. In study B (Graessner et al. 2021), right PGa did not distinguish pseudoword and real-word phrases.

These results are in line with a previous neuroimaging meta-analysis demonstrating a more consistent recruitment of left than right AG during semantic processing (Binder et al. 2009). However, they are inconsistent with meta-analyses suggesting exclusive recruitment of left but not right AG (Jackson 2021; Hodgson et al. 2021). Our findings suggest that right AG is also sensitive to semantics, but plays a weaker role than left AG, at least under “normal” conditions in young and healthy human adults. In support of this view, Jung-Beeman (2005) summarized evidence that both the left and right hemispheres are engaged in semantic cognition; however, the right hemisphere seems to perform coarser computations than the left.

As a hypothesis for future work, we propose that right AG might compensate when left AG is perturbed or even damaged. Such potential mechanisms of adaptive plasticity could be investigated in future studies combining non-invasive brain stimulation (e.g., TMS) with a neuroimaging readout (e.g., fMRI) (Bergmann et al. 2016; Hartwigsen and Volz 2021).

### Functional-Anatomical Subdivision

We observed distinct response profiles for the cytoarchitectonic subregions of the AG, *PGa* and *PGp*. Specifically, our combined analysis across all studies revealed that domain-general task difficulty effects were stronger in PGa than PGp, regardless of hemisphere.

These results partially support and partially refute previous functional-anatomical subdivisions of the AG. Noonan et al. (2013) performed a meta-analysis of functional neuroimaging studies on executive control during semantic processing. They found that left dorsal AG (~PGa) and adjacent intraparietal sulcus (IPS) showed increased activity for semantic tasks with a higher executive demand. In contrast, left ventral AG (~PGp) was engaged for semantic vs. phonological tasks, but insensitive to executive demands. Accordingly, the “controlled semantic cognition” (CSC) framework proposes that dorsal AG / IPS supports the controlled retrieval of semantic representations, rather than semantic representation *per se* (Jefferies 2013). Specifically, under this framework, left dorsal AG / IPS is associated with the multiple demand network (MDN) involved in domain-general executive control (Duncan 2010), whereas ventral AG supports semantic integration (Jefferies 2013). Similarly, Humphreys et al. (2021) argued that dorsal AG / IPS and ventral AG have distinct functions as dorsal AG shows a greater response when difficulty is increased, whereas ventral AG shows lower activity for harder tasks. In line with these proposals, we found stronger difficulty effects in PGa than PGp, suggesting a stronger contribution of PGa to domain-general task-difficulty-related processes. However, in contrast to these previous views, we found the same *negative* relationship between neural activity and task difficulty in all AG-ROIs: Activity decreased with increasing difficulty.

This response pattern contradicts the expected response pattern of a control-related MDN region (i.e., increased activity for increased difficulty) and is more consistent with the response of a DMN region (cf. Noonan et al. 2013). Previous reports of control-related activation in PGa may have reflected activation “spillover” from nearby MDN regions, such as IPS (Duncan 2010; Whitney et al. 2012). This is especially plausible in a coordinate-based meta-analysis (as in Noonan et al. 2013), which involves smoothing activation peaks in standard space using Gaussian kernels (Eickhoff et al. 2009). Indeed, a more recent meta-analysis of semantic control, which updated the Noonan et al. (2013) study with novel data and methodology, found no consistent AG engagement (Jackson 2021; also see Hodgson et al. 2021). Moreover, a high-resolution subject-specific parcellation study revealed that IPS is part of the core MDN, whereas the AG is anti-correlated with the MDN and more likely to belong to the DMN (Assem et al. 2020). Together with our finding of lower AG activity for harder tasks, these results oppose the view of the CSC framework that (part of) the AG is involved in executive control processes during semantic processing. Overall, if the semantic system is indeed composed of representation and control regions as the CSC framework proposes (Lambon Ralph et al. 2016), our results suggest that the AG supports semantic representation, rather than control. However, it is unclear whether representation and control can be strictly divided (Chapman et al. 2020), and there may be further subdivisions of the semantic system, such as long-term (semantic memory) vs. short-term (working memory) representation (Martin et al. 1994; Vigneau et al. 2006).

Crucially, in our study, all AG subregions—including bilateral PGa—also showed domain-specific semantic effects, which were separable from their domain-general task difficulty effects. These findings strongly suggest that bilateral PGa supports not only domain-general processes, but also domain-specific semantic processes (Mattheiss et al. 2018). Indeed, while the combined analysis across all studies indicated similar semantic effects (i.e., activity differences between semantic and non-semantic conditions) for PGa and PGp, left PGa seemed to exhibit the highest sensitivity to fine-grained semantic manipulations in individual studies. For example, in the study by Graessner et al. (2021), left PGa was the only AG region that showed an activity difference between meaningful and anomalous phrases when this subtle semantic difference was task-relevant. In Kuhnke et al. (2020b), left PGa was the only AG subregion that selectively responded to action and sound features of word meaning when these were task-relevant. These results suggest that left PGa might be the AG subregion that is most relevant for semantic cognition.

This view of left PGa is supported by Seghier (2013) who subdivided the left AG into anterior-dorsal and ventral subregions based on four functional neuroimaging studies (Sharp et al. 2009; Nelson et al. 2010; Seghier et al. 2010; Price and Ansari 2011). The anterior-dorsal subregion shows a remarkably close and consistent correspondence with area PGa (see Figure 4 in Seghier 2013). Moreover, in all four studies, this area was sensitive to semantic variables. The other (more ventral) subregions were more variable across studies, both in location and in relation to semantic processing. Taken together, these previous findings and our current results support the view that left PGa constitutes a functional unit that can be functionally distinguished from its neighbors (e.g., PGp, IPS) and that is engaged in semantic cognition.

### The Role of the AG in Semantics

A common view holds that the AG acts as a cross-modal convergence zone or “hub” that binds and integrates semantic features related to various sensory-motor modalities (Damasio 1989; Mesulam 1998; Binder and Desai 2011). This view is supported by the AG’s location at the junction between several sensory-motor processing streams (e.g., somatomotor, auditory, visual; Seghier 2013; Margulies et al. 2016). Moreover, the AG shows extensive structural (Hagmann et al. 2008; Bonner and Price 2013) and functional (Tomasi and Volkow 2011; Kuhnke et al. 2021) connectivity with various sensory-motor cortices. Crucially, functional neuroimaging studies indicate that AG activity increases with the amount of semantic information that can be extracted from a given input (Binder 2016). At the level of individual concepts, AG activity is modulated by thematic associations (Bar and Aminoff 2003), concreteness (Binder et al. 2005), frequency (Graves et al. 2010), and familiarity (Woodard et al. 2007). Beyond the single-concept level, the AG is sensitive to compositionality at the phrase (Price et al. 2015b), sentence (Obleser et al. 2007), and narrative (Ferstl et al. 2008) levels. Taken together, these results suggest that the AG integrates different semantic features into a coherent conceptual representation.

However, the role of a cross-modal hub of the semantic system is classically associated with the anterior temporal lobes (ATL) (Lambon Ralph et al. 2016). In support of this view, evidence from semantic dementia (Patterson et al. 2007; Jefferies 2013), functional neuroimaging (Visser et al. 2010; Rice et al. 2015), and TMS (Pobric et al. 2010a, b) indicates a crucial role of the ATL in semantic processing across virtually all types of concepts. If the ATL already acts as a cross-modal hub, what could be the function of the AG?

We propose that the AG constitutes a “multimodal” hub, whereas the ATL is an “amodal” hub. As an “amodal” hub, the ATL integrates semantic features into highly abstract representations that do not retain modality-specific information. As a “multimodal” hub, on the other hand, the AG binds different semantic features associated with the same concept, while retaining modality-specific information. During online processing, the AG could thereby enable efficient access to task-relevant semantic features (Kuhnke et al. 2020b, 2021).

Similar proposals have been put forth previously. For instance, Seghier (2013) argued that while integration and amodality have been associated with the ATL, the AG might support “first-order” integration that provides direct access to conceptual representations. Similarly, Reilly et al. (2016) proposed the AG to constitute a “low-order hub” engaged in multimodal feature binding, whereas the ATL acts as a “high-order” hub performing symbolic transformations on the bound features. During these non-linear transformations, modality-specific information is lost.

The multimodal–amodal hub theory is supported by several studies. Fernandino et al. (2016) found that AG activity during concreteness judgements correlated with the strength of sensory-motor associations for all modalities tested (action, sound, shape, color, motion). In contrast, ATL activity did not correlate with individual sensory-motor associations. In line with these results, Kuhnke et al. (2020b) demonstrated that left AG responds to both sound and action features of concepts when these are task-relevant. Again, the ATL did not show modality-specific effects. However, the ATL was engaged for abstract semantic information (i.e., words vs. pseudowords). In a follow-up study (Kuhnke et al. 2021), left AG was functionally coupled with auditory brain regions during sound feature retrieval, and with somatomotor regions during action feature retrieval. This suggests that left AG guides the retrieval of task-relevant semantic features via flexible coupling with different sensory-motor cortices. In contrast, the ATL interacted with other high-level cross-modal areas, but not sensory-motor regions. Finally, TMS over left AG can selectively disrupt the retrieval of individual task-relevant semantic features (Kuhnke et al. 2020a; also see Pobric et al. 2010a; Ishibashi et al. 2011). In contrast, TMS over ATL typically impairs semantic processing for all types of concepts (Pobric et al. 2010a, b).

The multimodal vs. amodal distinction may also explain the well-known thematic vs. taxonomic distinction between AG and ATL (Davis and Yee 2019). Establishing thematic relationships requires binding different elements associated with the same type of event, while retaining information about the individual elements (Mirman et al. 2017). For example, *banana* and *monkey* often co-occur, leading to a binding of these two concepts in a thematic association. Therefore, thematic relationships are “multimodal”. On the other hand, establishing taxonomic relationships often requires abstracting away from the individual concepts to an abstract similarity structure that transcends individual modalities (Lambon Ralph et al. 2010). Such an abstract representation seems necessary to explain the emergence of coherent semantic categories, which can combine superficially distinct entities (e.g., *pear* and *pineapple*) and distinguish superficially similar entities (e.g., *pear* and *lightbulb*). Thus, taxonomic relationships are “amodal”. Hence, the multimodal vs. amodal distinction can explain why thematic relationships are associated with the “multimodal” AG, whereas taxonomic relationships are associated with the “amodal” ATL (Schwartz et al. 2011; de Zubicaray et al. 2013). This view is also supported by study A (Chapman and Hartwigsen 2021), where AG activity was consistently higher for thematic than taxonomic and unrelated object pairs. Notably, however, AG activity was also relatively higher for taxonomic than unrelated pairs, suggesting that the thematic vs. taxonomic distinction is graded, rather than completely binary.

Overall, our and previous findings suggest that AG and ATL play distinct, complementary roles during semantic processing. ATL acts as an “amodal” hub that represents an abstract similarity structure transcending individual modalities (Patterson and Lambon Ralph 2016). In contrast, AG acts as a “multimodal” hub that binds different semantic features of the same concept, enabling efficient access to task-relevant features (Seghier 2013; Reilly et al. 2016; Kuhnke et al. 2020b, 2021).

## Conclusion

In conclusion, we found that AG responses are best explained by separable effects of task difficulty and semantic processing demand. Task difficulty effects were stronger in PGa than PGp, regardless of hemisphere. Semantic effects were stronger in left than right AG, regardless of subregion. These results indicate that the AG is involved in both domain-general task-difficulty-related processes and domain-specific semantic processes. In semantic processing, we propose that left AG acts as a “multimodal” hub which binds different semantic features associated with the same concept, enabling efficient retrieval of task-relevant features. While the right AG seems to play a weaker role under “normal” conditions in young and healthy adults, it is also sensitive to semantics and might compensate when the left AG is damaged.

## Statements & Declarations

## Acknowledgements

We wish to thank the medical technical assistants of the Max Planck Institute for Human Cognitive and Brain Sciences for their tremendous help during participant acquisition and fMRI measurements. Moreover, we thank two anonymous reviewers for their insightful comments on a previous version of this manuscript.

## Author Contributions

**Philipp Kuhnke**: Conceptualization, Investigation, Data curation, Formal analysis, Methodology, Writing—original draft, Writing—review and editing;

**Curtiss A. Chapman**: Conceptualization, Investigation, Data curation, Formal analysis, Methodology, Writing—original draft, Writing—review and editing;

**Vincent K.M. Cheung**: Conceptualization, Formal analysis, Methodology, Writing—review and editing;

**Sabrina Turker**: Investigation, Data curation;

**Astrid Graessner**: Investigation, Data curation;

**Sandra Martin**: Investigation, Data curation;

**Kathleen A. Williams**: Resources, Methodology;

**Gesa Hartwigsen**: Conceptualization, Funding acquisition, Supervision, Project administration, Writing—review and editing.

## References

Andrews-Hanna JR (2012) The Brain’s Default Network and Its Adaptive Role in Internal Mentation. Neurosci 18:251–270. https://doi.org/10.1177/1073858411403316

Assem M, Glasser MF, Van Essen DC, Duncan J (2020) A Domain-General Cognitive Core Defined in Multimodally Parcellated Human Cortex. Cereb Cortex 30:4361–4380. https://doi.org/10.1093/cercor/bhaa023

Bar M, Aminoff E (2003) Cortical Analysis of Visual Context. Neuron 38:347–358. https://doi.org/10.1016/S0896-6273(03)00167-3

Bassett DS, Sporns O (2017) Network neuroscience. Nat Neurosci 20:353–364. https://doi.org/10.1038/nn.4502

Bates D, Mächler M, Bolker B, Walker S (2015) Fitting Linear Mixed-Effects Models Using lme4. J Stat Softw 67:1–48. https://doi.org/10.18637/jss.v067.i01

Bergmann TO, Karabanov A, Hartwigsen G, et al (2016) Combining non-invasive transcranial brain stimulation with neuroimaging and electrophysiology: Current approaches and future perspectives. Neuroimage 140:4–19. https://doi.org/10.1016/j.neuroimage.2016.02.012

Binder JR (2016) In defense of abstract conceptual representations. Psychon Bull Rev 23:1096–1108. https://doi.org/10.3758/s13423-015-0909-1

Binder JR, Desai RH (2011) The neurobiology of semantic memory. Trends Cogn Sci 15:527– 536. https://doi.org/10.1016/j.tics.2011.10.001

Binder JR, Desai RH, Graves WW, Conant LL (2009) Where Is the Semantic System? A Critical Review and Meta-Analysis of 120 Functional Neuroimaging Studies. Cereb Cortex 19:2767–2796. https://doi.org/10.1093/cercor/bhp055

Binder JR, Fernandino L (2015) Semantic Processing. In: Brain Mapping. Elsevier, Amsterdam, pp 445–454

Binder JR, Frost JA, Hammeke TA, et al (1999) Conceptual Processing during the Conscious Resting State: A Functional MRI Study. J Cogn Neurosci 11:80–93.https://doi.org/10.1162/089892999563265

Binder JR, Westbury CF, McKiernan KA, et al (2005) Distinct Brain Systems for Processing Concrete and Abstract Concepts. J Cogn Neurosci 17:905–917. https://doi.org/10.1162/0898929054021102

Bolker B (2020) Maximum likelihood estimation and analysis with the bbmle package. Citeseer

Bonner MF, Peelle JE, Cook PA, Grossman M (2013) Heteromodal conceptual processing in the angular gyrus. Neuroimage 71:175–186. https://doi.org/10.1016/j.neuroimage.2013.01.006

Bonner MF, Price AR (2013) Where Is the Anterior Temporal Lobe and What Does It Do? J Neurosci 33:4213–4215. https://doi.org/10.1523/JNEUROSCI.0041-13.2013

Brett M, Anton J-L, Valabregue R, Poline J-B (2002) Region of interest analysis using an SPM toolbox [abstract]. Sendai, Japan

Buckner RL, Andrews-Hanna JR, Schacter DL (2008) The brain’s default network: Anatomy, function, and relevance to disease. Ann N Y Acad Sci 1124:1–38. https://doi.org/10.1196/annals.1440.011

Burnham KP, Anderson DR (2004) Multimodel Inference: Understanding AIC and BIC in Model Selection. Sociol Methods Res 33:261–304. https://doi.org/10.1177/0049124104268644

Caspers S, Eickhoff SB, Geyer S, et al (2008) The human inferior parietal lobule in stereotaxic space. Brain Struct Funct 212:481–495. https://doi.org/10.1007/s00429-008-0195-z

Caspers S, Geyer S, Schleicher A, et al (2006) The human inferior parietal cortex: Cytoarchitectonic parcellation and interindividual variability. Neuroimage 33:430–448. https://doi.org/10.1016/j.neuroimage.2006.06.054

Chai LR, Mattar MG, Blank IA, et al (2016) Functional Network Dynamics of the Language System. Cereb Cortex 26:4148–4159. https://doi.org/10.1093/cercor/bhw238

Chapman CA, Hartwigsen G (2021) Semantic conflict is resolved by semantic and multiple demand networks. In: Poster presented at the 13th Meeting of the Society for the Neurobiology of Language, October 5–8, 2021 (virtual edition).

Chapman CA, Hasan O, Schulz PE, Martin RC (2020) Evaluating the distinction between semantic knowledge and semantic access: Evidence from semantic dementia and comprehension-impaired stroke aphasia. Psychon Bull Rev 27:607–639.https://doi.org/10.3758/s13423-019-01706-6

Damasio AR (1989) The Brain Binds Entities and Events by Multiregional Activation from Convergence Zones. Neural Comput 1:123–132. https://doi.org/10.1162/neco.1989.1.1.123

Davey J, Cornelissen PL, Thompson HE, et al (2015) Automatic and Controlled Semantic Retrieval: TMS Reveals Distinct Contributions of Posterior Middle Temporal Gyrus and Angular Gyrus. J Neurosci 35:15230–15239. https://doi.org/10.1523/JNEUROSCI.4705-14.2015

Davis CP, Yee E (2019) Features, labels, space, and time: factors supporting taxonomic relationships in the anterior temporal lobe and thematic relationships in the angular gyrus. Lang Cogn Neurosci 34:1347–1357. https://doi.org/10.1080/23273798.2018.1479530

de Zubicaray GI, Hansen S, McMahon KL (2013) Differential processing of thematic and categorical conceptual relations in spoken word production. J Exp Psychol Gen 142:131– 142. https://doi.org/10.1037/a0028717

Desai RH, Reilly M, van Dam W (2018) The multifaceted abstract brain. Philos Trans R Soc B Biol Sci 373:20170122. https://doi.org/10.1098/rstb.2017.0122

Duffau H, Capelle L, Sichez JP, et al (1999) Intra-Operative Direct Electrical Stimulations of the Central Nervous System: The Salpêtrière Experience With 60 Patients. Acta Neurochir (Wien) 141:1157–1167. https://doi.org/10.1007/s007010050413

Duffau H, Velut S, Mitchell M-C, et al (2004) Intra-operative mapping of the subcortical visual pathways using direct electrical stimulations. Acta Neurochir (Wien) 146:265–270. https://doi.org/10.1007/s00701-003-0199-7

Duncan J (2010) The multiple-demand (MD) system of the primate brain: mental programs for intelligent behaviour. Trends Cogn Sci 14:172–179. https://doi.org/10.1016/j.tics.2010.01.004

Eickhoff SB, Heim S, Zilles K, Amunts K (2006) Testing anatomically specified hypotheses in functional imaging using cytoarchitectonic maps. Neuroimage 32:570–582. https://doi.org/10.1016/j.neuroimage.2006.04.204

Eickhoff SB, Laird AR, Grefkes C, et al (2009) Coordinate-based activation likelihood estimation meta-analysis of neuroimaging data: A random-effects approach based on empirical estimates of spatial uncertainty. Hum Brain Mapp 30:2907–2926. https://doi.org/10.1002/hbm.20718

Eickhoff SB, Stephan KE, Mohlberg H, et al (2005) A new SPM toolbox for combining probabilistic cytoarchitectonic maps and functional imaging data. Neuroimage 25:1325– 1335. https://doi.org/10.1016/j.neuroimage.2004.12.034

Fedorenko E, Kanwisher N (2009) Neuroimaging of Language: Why Hasn’t a Clearer Picture Emerged? Lang Linguist Compass 3:839–865. https://doi.org/10.1111/j.1749-818X.2009.00143.x

Fernandino L, Binder JR, Desai RH, et al (2016) Concept Representation Reflects Multimodal Abstraction: A Framework for Embodied Semantics. Cereb Cortex 26:2018–2034. https://doi.org/10.1093/cercor/bhv020

Ferstl EC, Neumann J, Bogler C, von Cramon DY (2008) The extended language network: A meta-analysis of neuroimaging studies on text comprehension. Hum Brain Mapp 29:581– 593. https://doi.org/10.1002/hbm.20422

Finn ES (2021) Is it time to put rest to rest? Trends Cogn Sci 25:1021–1032. https://doi.org/10.1016/j.tics.2021.09.005

Graessner A, Zaccarella E, Hartwigsen G (2021) Differential contributions of left-hemispheric language regions to basic semantic composition. Brain Struct Funct 226:501–518. https://doi.org/10.1007/s00429-020-02196-2

Graves WW, Binder JR, Desai RH, et al (2010) Neural correlates of implicit and explicit combinatorial semantic processing. Neuroimage 53:638–646. https://doi.org/10.1016/j.neuroimage.2010.06.055

Hagmann P, Cammoun L, Gigandet X, et al (2008) Mapping the Structural Core of Human Cerebral Cortex. PLoS Biol 6:e159. https://doi.org/10.1371/journal.pbio.0060159

Hahn B, Ross TJ, Yang Y, et al (2007) Nicotine Enhances Visuospatial Attention by Deactivating Areas of the Resting Brain Default Network. J Neurosci 27:3477–3489. https://doi.org/10.1523/JNEUROSCI.5129-06.2007

Hartwigsen G, Volz LJ (2021) Probing rapid network reorganization of motor and language functions via neuromodulation and neuroimaging. Neuroimage 224:117449. https://doi.org/10.1016/j.neuroimage.2020.117449

Hartwigsen G, Weigel A, Schuschan P, et al (2016) Dissociating Parieto-Frontal Networks for Phonological and Semantic Word Decisions: A Condition-and-Perturb TMS Study. Cereb Cortex 26:2590–2601. https://doi.org/10.1093/cercor/bhv092

Hauk O, Tschentscher N (2013) The Body of Evidence: What Can Neuroscience Tell Us about Embodied Semantics? Front Psychol 4:1–14. https://doi.org/10.3389/fpsyg.2013.00050

Hodgson VJ, Lambon Ralph MA, Jackson RL (2021) Multiple dimensions underlying the functional organization of the language network. Neuroimage 241:118444. https://doi.org/10.1016/j.neuroimage.2021.118444

Hoenig K, Sim E-J, Bochev V, et al (2008) Conceptual Flexibility in the Human Brain: Dynamic Recruitment of Semantic Maps from Visual, Motor, and Motion-related Areas. J Cogn Neurosci 20:1799–1814. https://doi.org/10.1162/jocn.2008.20123

Humphreys GF, Hoffman P, Visser M, et al (2015) Establishing task-and modality-dependent dissociations between the semantic and default mode networks. Proc Natl Acad Sci 112:7857–7862. https://doi.org/10.1073/pnas.1422760112

Humphreys GF, Lambon Ralph MA (2017) Mapping domain-selective and counterpointed domain-general higher cognitive functions in the lateral parietal cortex: Evidence from fMRI comparisons of difficulty-varying semantic versus visuo-spatial tasks, and functional connectivity analyses. Cereb Cortex 27:4199–4212. https://doi.org/10.1093/cercor/bhx107

Humphreys GF, Lambon Ralph MA, Simons JS (2021) A Unifying Account of Angular Gyrus Contributions to Episodic and Semantic Cognition. Trends Neurosci 44:452–463. https://doi.org/10.1016/j.tins.2021.01.006

Ishibashi R, Lambon Ralph MA, Saito S, Pobric G (2011) Different roles of lateral anterior temporal lobe and inferior parietal lobule in coding function and manipulation tool knowledge: Evidence from an rTMS study. Neuropsychologia 49:1128–1135. https://doi.org/10.1016/j.neuropsychologia.2011.01.004

Jackson RL (2021) The neural correlates of semantic control revisited. Neuroimage 224:. https://doi.org/10.1016/j.neuroimage.2020.117444

Jefferies E (2013) The neural basis of semantic cognition: Converging evidence from neuropsychology, neuroimaging and TMS. Cortex 49:611–625. https://doi.org/10.1016/j.cortex.2012.10.008

Jung-Beeman M (2005) Bilateral brain processes for comprehending natural language. Trends Cogn Sci 9:512–518. https://doi.org/10.1016/j.tics.2005.09.009

Kemmerer D (2015) Are the motor features of verb meanings represented in the precentral motor cortices? Yes, but within the context of a flexible, multilevel architecture for conceptual knowledge. Psychon Bull Rev 22:1068–1075. https://doi.org/10.3758/s13423-014-0784-1

Kiefer M, Pulvermüller F (2012) Conceptual representations in mind and brain: Theoretical developments, current evidence and future directions. Cortex 48:805–825. https://doi.org/10.1016/j.cortex.2011.04.006

Kuhnke P, Beaupain MC, Cheung VKM, et al (2020a) Left posterior inferior parietal cortex causally supports the retrieval of action knowledge. Neuroimage 219:117041. https://doi.org/10.1016/j.neuroimage.2020.117041

Kuhnke P, Kiefer M, Hartwigsen G (2020b) Task-Dependent Recruitment of Modality-Specific and Multimodal Regions during Conceptual Processing. Cereb Cortex 30:3938–3959. https://doi.org/10.1093/cercor/bhaa010

Kuhnke P, Kiefer M, Hartwigsen G (2021) Task-Dependent Functional and Effective Connectivity during Conceptual Processing. Cereb Cortex 31:3475–3493. https://doi.org/10.1093/cercor/bhab026

Kuznetsova A, Brockhoff PB, Christensen RHB (2017) lmerTest Package: Tests in Linear Mixed Effects Models. J Stat Softw 82:. https://doi.org/10.18637/jss.v082.i13

Lambon Ralph MA (2014) Neurocognitive insights on conceptual knowledge and its breakdown.Philos Trans R Soc B Biol Sci 369:20120392. https://doi.org/10.1098/rstb.2012.0392

Lambon Ralph MA, Jefferies E, Patterson K, Rogers TT (2016) The neural and computational bases of semantic cognition. Nat Rev Neurosci 18:42–55.https://doi.org/10.1038/nrn.2016.150

Lambon Ralph MA, Sage K, Jones RW, Mayberry EJ (2010) Coherent concepts are computed in the anterior temporal lobes. Proc Natl Acad Sci 107:2717–2722.https://doi.org/10.1073/pnas.0907307107

Lüdecke D (2021) sjPlot: Data visualization for statistics in social science. Zenodo. https://doi.org/10.5281/zenodo.2400856

Lüdecke D (2018) ggeffects: Tidy Data Frames of Marginal Effects from Regression Models. J Open Source Softw 3:772. https://doi.org/10.21105/joss.00772

Margulies DS, Ghosh SS, Goulas A, et al (2016) Situating the default-mode network along a principal gradient of macroscale cortical organization. Proc Natl Acad Sci 113:12574– 12579. https://doi.org/10.1073/pnas.1608282113

Martin RC, Shelton JR, Yaffee LS (1994) Language Processing and Working Memory: Neuropsychological Evidence for Separate Phonological and Semantic Capacities. J Mem Lang 33:83–111. https://doi.org/10.1006/jmla.1994.1005

Martin S, Saur D, Hartwigsen G (2021) Age-Dependent Contribution of Domain-General Networks to Semantic Cognition. Cereb Cortex 1–57. https://doi.org/10.1093/cercor/bhab252

Mattheiss SR, Levinson H, Graves WW (2018) Duality of Function: Activation for Meaningless Nonwords and Semantic Codes in the Same Brain Areas. Cereb Cortex 28:2516–2524. https://doi.org/10.1093/cercor/bhy053

Mesulam MM (1998) From sensation to cognition. Brain 121:1013–1052. https://doi.org/10.1093/brain/121.6.1013

Mirman D, Landrigan J-F, Britt AE (2017) Taxonomic and thematic semantic systems. Psychol Bull 143:499–520. https://doi.org/10.1037/bul0000092

Morcom AM, Fletcher PC (2007) Does the brain have a baseline? Why we should be resisting a rest. Neuroimage 37:1073–1082. https://doi.org/10.1016/j.neuroimage.2006.09.013

Nelson SM, Cohen AL, Power JD, et al (2010) A Parcellation Scheme for Human Left Lateral Parietal Cortex. Neuron 67:156–170. https://doi.org/10.1016/j.neuron.2010.05.025

Noonan KA, Jefferies E, Visser M, Lambon Ralph MA (2013) Going beyond Inferior Prefrontal Involvement in Semantic Control: Evidence for the Additional Contribution of Dorsal Angular Gyrus and Posterior Middle Temporal Cortex. J Cogn Neurosci 25:1824–1850. https://doi.org/10.1162/jocn_a_00442

Obleser J, Wise RJS, Dresner MA, Scott SK (2007) Functional integration across brain regions improves speech perception under adverse listening conditions. J Neurosci 27:2283–2289. https://doi.org/10.1523/JNEUROSCI.4663-06.2007

Patterson K, Lambon Ralph MA (2016) The hub-and-spoke hypothesis of semantic memory. In: Neurobiology of language. Elsevier, Amsterdam, pp 765–775

Patterson K, Nestor PJ, Rogers TT (2007) Where do you know what you know? The representation of semantic knowledge in the human brain. Nat Rev Neurosci 8:976–987. https://doi.org/10.1038/nrn2277

Pobric G, Jefferies E, Lambon Ralph MA (2010a) Category-Specific versus Category-General Semantic Impairment Induced by Transcranial Magnetic Stimulation. Curr Biol 20:964–968. https://doi.org/10.1016/j.cub.2010.03.070

Pobric G, Jefferies E, Lambon Ralph MA (2010b) Amodal semantic representations depend on both anterior temporal lobes: Evidence from repetitive transcranial magnetic stimulation. Neuropsychologia 48:1336–1342. https://doi.org/10.1016/j.neuropsychologia.2009.12.036

Price AR, Bonner MF, Grossman M (2015a) Semantic Memory: Cognitive and Neuroanatomical Perspectives. In: Brain Mapping. Elsevier, Amsterdam, pp 529–536

Price AR, Bonner MF, Peelle JE, Grossman M (2015b) Converging Evidence for the Neuroanatomic Basis of Combinatorial Semantics in the Angular Gyrus. J Neurosci 35:3276–3284. https://doi.org/10.1523/JNEUROSCI.3446-14.2015

Price AR, Peelle JE, Bonner MF, et al (2016) Causal Evidence for a Mechanism of Semantic Integration in the Angular Gyrus as Revealed by High-Definition Transcranial Direct Current Stimulation. J Neurosci 36:3829–3838. https://doi.org/10.1523/JNEUROSCI.3120-15.2016

Price GR, Ansari D (2011) Symbol processing in the left angular gyrus: Evidence from passive perception of digits. Neuroimage 57:1205–1211. https://doi.org/10.1016/j.neuroimage.2011.05.035

Rabe M, Vasishth S, Hohenstein S, et al (2020) hypr: An R package for hypothesis-driven contrast coding. J Open Source Softw 5:2134. https://doi.org/10.21105/joss.02134

Raichle ME (2015) The Brain’s Default Mode Network. Annu Rev Neurosci 38:433–447. https://doi.org/10.1146/annurev-neuro-071013-014030

Reilly J, Peelle JE, Garcia A, Crutch SJ (2016) Linking somatic and symbolic representation in semantic memory: the dynamic multilevel reactivation framework. Psychon Bull Rev 23:1002–1014. https://doi.org/10.3758/s13423-015-0824-5

Rice GE, Lambon Ralph MA, Hoffman P (2015) The Roles of Left Versus Right Anterior Temporal Lobes in Conceptual Knowledge: An ALE Meta-analysis of 97 Functional Neuroimaging Studies. Cereb Cortex 25:4374–4391. https://doi.org/10.1093/cercor/bhv024

Schad DJ, Vasishth S, Hohenstein S, Kliegl R (2020) How to capitalize on a priori contrasts in linear (mixed) models: A tutorial. J Mem Lang 110:104038. https://doi.org/10.1016/j.jml.2019.104038

Schwartz MF, Kimberg DY, Walker GM, et al (2011) Neuroanatomical dissociation for taxonomic and thematic knowledge in the human brain. Proc Natl Acad Sci 108:8520– 8524. https://doi.org/10.1073/pnas.1014935108

Seghier ML (2013) The angular gyrus: Multiple functions and multiple subdivisions. Neuroscientist 19:43–61. https://doi.org/10.1177/1073858412440596

Seghier ML, Fagan E, Price CJ (2010) Functional Subdivisions in the Left Angular Gyrus Where the Semantic System Meets and Diverges from the Default Network. J Neurosci 30:16809– 16817. https://doi.org/10.1523/JNEUROSCI.3377-10.2010

Sharp DJ, Awad M, Warren JE, et al (2009) The neural response to changing semantic and perceptual complexity during language processing. Hum Brain Mapp 31:365–377. https://doi.org/10.1002/hbm.20871

Sliwinska MW, James A, Devlin JT (2015) Inferior Parietal Lobule Contributions to Visual Word Recognition. J Cogn Neurosci 27:593–604. https://doi.org/10.1162/jocn_a_00721

Stark CEL, Squire LR (2001) When zero is not zero: The problem of ambiguous baseline conditions in fMRI. Proc Natl Acad Sci U S A 98:12760–12765.https://doi.org/10.1073/pnas.221462998

Tomasi D, Volkow ND (2011) Association between Functional Connectivity Hubs and Brain Networks. Cereb Cortex 21:2003–2013. https://doi.org/10.1093/cercor/bhq268

Turker S, Kuhnke P, Hartwigsen G (2021) The role of the left temporo-parietal cortex for pseudoword processing: evidence from combined neuroimaging and brain stimulation. In: Abstract accepted for the 19th Old World Conference on Phonology - Workshop: Phonology & Dyslexia.

van Elk M, van Schie H, Bekkering H (2014) Action semantics: A unifying conceptual framework for the selective use of multimodal and modality-specific object knowledge. Phys Life Rev 11:220–250. https://doi.org/10.1016/j.plrev.2013.11.005

Vigneau M, Beaucousin V, Hervé PY, et al (2006) Meta-analyzing left hemisphere language areas: Phonology, semantics, and sentence processing. Neuroimage 30:1414–1432. https://doi.org/10.1016/j.neuroimage.2005.11.002

Visser M, Jefferies E, Lambon Ralph MA (2010) Semantic Processing in the Anterior Temporal Lobes: A Meta-analysis of the Functional Neuroimaging Literature. J Cogn Neurosci 22:1083–1094. https://doi.org/10.1162/jocn.2009.21309

Whitney C, Kirk M, O’Sullivan J, et al (2012) Executive Semantic Processing Is Underpinned by a Large-scale Neural Network: Revealing the Contribution of Left Prefrontal, Posterior Temporal, and Parietal Cortex to Controlled Retrieval and Selection Using TMS. J Cogn Neurosci 24:133–147. https://doi.org/10.1162/jocn_a_00123

Woodard JL, Seidenberg M, Nielson KA, et al (2007) Temporally Graded Activation of Neocortical Regions in Response to Memories of Different Ages. J Cogn Neurosci 19:1113–1124. https://doi.org/10.1162/jocn.2007.19.7.1113

