## Supplementary Material for "The role of the angular gyrus in semantic cognition – A synthesis of five functional neuroimaging studies"

(Kuhnke, Chapman et al. 2022 BSF)

### Supplementary Results

#### Behavioral analyses

The following tables report the results of statistical analyses on the behavioral measures, i.e. error rates and mean response times for correct trials. For statistical inference, we first conducted repeated-measures ANOVAs. Significant interactions were resolved using post-hoc paired t-tests. Bold font highlights significant effects ($p < 0.05$ Bonferroni-Holm corrected for multiple comparisons); italic fonts highlights trends ($p < 0.05$ uncorrected).

**Table S1**. Study A (Chapman and Hartwigsen 2021): Repeated-measures ANOVA.

| Effect | df | F | p | partial $\eta^2$ |
|---|---|---|---|---|
| **Error Rates** | | | | |
| TASK | 1,39 | 8.595 | **0.006** | 0.181 |
| CONDITION | 2,78 | 13.935 | **<0.001** | 0.263 |
| TASK x CONDITION | 2,78 | 8.881 | **<0.001** | 0.185 |
| **Mean RTs** | | | | |
| TASK | 1,39 | 11.652 | **0.002** | 0.230 |
| CONDITION | 2,78 | 9.853 | **<0.001** | 0.202 |
| TASK x CONDITION | 2,78 | 10.293 | **<0.001** | 0.209 |

**Table S2**. Study A (Chapman and Hartwigsen 2021): Post-hoc paired t-tests.

| Comparison | Mean | SE | T | p (unc.) | p (corr.) |
|---|---|---|---|---|---|
| **Error Rates** | | | | | |
| Taxonomic vs. thematic task | -2.219 | 0.757 | -2.932 | **0.006** | **0.042** |
| Taxonomic vs. scrambled task | -0.722 | 0.653 | -1.106 | 0.276 | 0.729 |
| Thematic vs. scrambled task | 1.496 | 0.772 | 1.938 | 0.060 | 0.240 |
| Taxonomic task: Taxonomic vs. thematic | 3.578 | 1.150 | 3.111 | **0.003** | **0.024** |
| Taxonomic task: Taxonomic vs. unrelated | 6.995 | 1.194 | 5.860 | **<0.001** | **<0.001** |
| Taxonomic task: Thematic vs. unrelated | 3.417 | 0.662 | 5.165 | **<0.001** | **<0.001** |
| Thematic task: Taxonomic vs. thematic | -3.110 | 1.275 | -2.440 | *0.019* | *0.095* |
| Thematic task: Taxonomic vs. unrelated | 0.963 | 1.349 | 0.714 | 0.479 | 0.552 |
| Thematic task: Thematic vs. unrelated | 4.073 | 1.501 | 2.713 | *0.010* | *0.060* |

| | | | | | |
|---|---|---|---|---|---|
| Scrambled task: Congruent vs. incongruent | 0.990 | 0.834 | 1.187 | 0.243 | 0.729 |
| **Mean RTs** | | | | | |
| Taxonomic vs. thematic task | -82.321 | 24.117 | -3.413 | **0.002** | **0.010** |
| Taxonomic vs. scrambled task | -350.935 | 39.198 | -8.953 | **<0.001** | **<0.001** |
| Thematic vs. scrambled task | -268.613 | 43.922 | -6.116 | **<0.001** | **<0.001** |
| Taxonomic task: Taxonomic vs. thematic | -48.785 | 25.632 | -1.903 | 0.064 | 0.256 |
| Taxonomic task: Taxonomic vs. unrelated | 101.348 | 25.950 | 3.905 | **<0.001** | **0.002** |
| Taxonomic task: Thematic vs. unrelated | 150.134 | 20.793 | 7.220 | **<0.001** | **<0.001** |
| Thematic task: Taxonomic vs. thematic | 0.065 | 27.350 | 0.002 | 0.998 | 1.000 |
| Thematic task: Taxonomic vs. unrelated | -10.509 | 24.008 | -0.438 | 0.664 | 1.000 |
| Thematic task: Thematic vs. unrelated | -10.574 | 20.461 | -0.517 | 0.608 | 1.000 |
| Scrambled task: Congruent vs. incongruent | 276.798 | 45.196 | 6.124 | **<0.001** | **<0.001** |

**Table S3**. Study B (Graessner et al. 2021): Repeated-measures ANOVA.

| **Effect** | **df** | **F** | **p** | **partial η$^2$** |
|---|---|---|---|---|
| **Error Rates** | | | | |
| TASK | 1,32 | 4.668 | **0.038** | 0.127 |
| CONDITION | 3,96 | 19.856 | **<0.001** | 0.383 |
| TASK x CONDITION | 3,96 | 24.287 | **<0.001** | 0.431 |
| **Mean RTs** | | | | |
| TASK | 1,32 | 10.462 | **0.003** | 0.246 |
| CONDITION | 3,96 | 46.066 | **<0.001** | 0.590 |
| TASK x CONDITION | 3,96 | 61.417 | **<0.001** | 0.657 |

**Table S4**. Study B (Graessner et al. 2021): Post-hoc paired t-tests.

| **Comparison** | **Mean** | **SE** | **T** | **p (unc.)** | **p (corr.)** |
|---|---|---|---|---|---|
| **Error Rates** | | | | | |
| Implicit task: Meaningful vs. anomalous | -5.195 | 0.940 | -5.525 | **<0.001** | **<0.001** |
| Implicit task: Meaningful vs. pseudoword | -6.223 | 0.808 | -7.701 | **<0.001** | **<0.001** |
| Implicit task: Meaningful vs. single words | 1.677 | 0.527 | 3.185 | **0.003** | **0.021** |
| Implicit task: Anomalous vs. pseudoword | -1.028 | 1.096 | -0.938 | 0.355 | 1.000 |
| Implicit task: Anomalous vs. single words | 6.872 | 1.008 | 6.820 | **<0.001** | **<0.001** |
| Implicit task: Pseudoword vs. single words | 7.900 | 0.920 | 8.591 | **<0.001** | **<0.001** |
| Explicit task: Meaningful vs. anomalous | 0.487 | 1.097 | 0.444 | 0.660 | 1.000 |
| Explicit task: Meaningful vs. pseudoword | 2.435 | 0.840 | 2.899 | **0.007** | **0.035** |
| Explicit task: Meaningful vs. single words | 2.922 | 0.899 | 3.250 | **0.003** | **0.021** |
| Explicit task: Anomalous vs. pseudoword | 1.948 | 0.908 | 2.147 | *0.040* | *0.160* |
| Explicit task: Anomalous vs. single words | 2.435 | 0.663 | 3.672 | **0.001** | **0.008** |
| Explicit task: Pseudoword vs. single words | 0.487 | 0.726 | 0.671 | 0.507 | 1.000 |
| **Mean RTs** | | | | | |
| Implicit task: Meaningful vs. anomalous | -73.727 | 7.755 | -9.507 | **<0.001** | **<0.001** |
| Implicit task: Meaningful vs. pseudoword | -208.421 | 14.163 | -14.716 | **<0.001** | **<0.001** |
| Implicit task: Meaningful vs. single words | -186.626 | 18.860 | -9.895 | **<0.001** | **<0.001** |

| | | | | | |
|---|---|---|---|---|---|
| Implicit task: Anomalous vs. pseudoword | -134.694 | 14.411 | -9.346 | **<0.001** | **<0.001** |
| Implicit task: Anomalous vs. single words | -112.899 | 20.350 | -5.548 | **<0.001** | **<0.001** |
| Implicit task: Pseudoword vs. single words | 21.795 | 16.876 | 1.291 | 0.206 | 0.206 |
| Explicit task: Meaningful vs. anomalous | -72.085 | 14.375 | -5.015 | **<0.001** | **<0.001** |
| Explicit task: Meaningful vs. pseudoword | 11.511 | 13.749 | 0.837 | 0.409 | 0.409 |
| Explicit task: Meaningful vs. single words | -64.077 | 13.898 | -4.610 | **<0.001** | **<0.001** |
| Explicit task: Anomalous vs. pseudoword | 83.595 | 12.563 | 6.654 | **<0.001** | **<0.001** |
| Explicit task: Anomalous vs. single words | 8.008 | 14.153 | 0.566 | 0.575 | 0.575 |
| Explicit task: Pseudoword vs. single words | -75.588 | 11.968 | -6.316 | **<0.001** | **<0.001** |

**Table S5**. Study C (Kuhnke et al. 2020): Repeated-measures ANOVA.

| **Effect** | **df** | **F** | **p** | **partial $\eta^2$** |
|---|---|---|---|---|
| **Error Rates** | | | | |
| TASK | 2,78 | 58.110 | **<0.001** | **0.598** |
| SOUND | 1,39 | 10.779 | **0.002** | **0.217** |
| ACTION | 1,39 | 16.109 | **<0.001** | **0.292** |
| TASK x SOUND | 2,78 | 40.548 | **<0.001** | **0.510** |
| TASK x ACTION | 2,78 | 8.432 | **0.003** | **0.178** |
| SOUND x ACTION | 1,39 | 112.272 | **<0.001** | **0.742** |
| TASK x SOUND x ACTION | 2,78 | 35.047 | **<0.001** | **0.473** |
| **Mean RTs** | | | | |
| TASK | 2,78 | 148.434 | **<0.001** | **0.792** |
| SOUND | 1,39 | 6.550 | **0.014** | **0.144** |
| ACTION | 1,39 | 26.038 | **<0.001** | **0.400** |
| TASK x SOUND | 2,78 | 0.333 | 0.678 | 0.008 |
| TASK x ACTION | 2,78 | 7.761 | **0.004** | **0.166** |
| SOUND x ACTION | 1,39 | 169.427 | **<0.001** | **0.813** |
| TASK x SOUND x ACTION | 2,78 | 71.260 | **<0.001** | **0.646** |

**Table S6**. Study C (Kuhnke et al. 2020): Post-hoc paired t-tests.

| **Comparison** | **Mean** | **SE** | **T** | **p (unc.)** | **p (corr.)** |
|---|---|---|---|---|---|
| **Error Rates** | | | | | |
| Lexical decisions: Words vs. Pseudowords | 0.625 | 0.356 | 1.756 | 0.087 | 0.783 |
| Lexical decisions: High sound-high action vs. high sound-low action | 0.000 | 0.684 | 0.000 | 1.000 | 1.000 |
| Lexical decisions: High sound-high action vs. low sound-high action | -0.417 | 0.512 | -0.813 | 0.421 | 1.000 |
| Lexical decisions: High sound-high action vs. low sound-low action | -0.729 | 0.475 | -1.535 | 0.133 | 1.000 |
| Lexical decisions: High sound-low action vs. low sound-high action | -0.417 | 0.616 | -0.676 | 0.503 | 1.000 |
| Lexical decisions: High sound-low action vs. low sound-low action | -0.729 | 0.729 | -1.000 | 0.323 | 1.000 |
| Lexical decisions: Low sound-high action vs. low sound-low action | -0.313 | 0.665 | -0.470 | 0.641 | 1.000 |
| Sound judgments: High sound-high action vs. high sound-low action | -19.375 | 2.093 | -9.257 | **<0.001** | **<0.001** |
| Sound judgments: High sound-high action vs. low sound-high action | -1.406 | 1.110 | -1.267 | 0.213 | 1.000 |

| | | | | | |
|---|---|---|---|---|---|
| Sound judgments: High sound-high action vs. low sound-low action | 2.292 | 0.934 | 2.453 | *0.019* | *0.190* |
| Sound judgments: High sound-low action vs. low sound-high action | 17.969 | 2.688 | 6.684 | **<0.001** | **<0.001** |
| Sound judgments: High sound-low action vs. low sound-low action | 21.667 | 2.750 | 7.880 | **<0.001** | **<0.001** |
| Sound judgments: Low sound-high action vs. low sound-low action | 3.698 | 0.788 | 4.695 | **<0.001** | **<0.001** |
| Action judgments: High sound-high action vs. high sound-low action | -10.104 | 1.952 | -5.177 | **<0.001** | **<0.001** |
| Action judgments: High sound-high action vs. low sound-high action | -12.604 | 1.663 | -7.580 | **<0.001** | **<0.001** |
| Action judgments: High sound-high action vs. low sound-low action | -4.896 | 1.922 | -2.548 | *0.015* | *0.165* |
| Action judgments: High sound-low action vs. low sound-high action | -2.500 | 2.579 | -0.969 | 0.338 | 1.000 |
| Action judgments: High sound-low action vs. low sound-low action | 5.208 | 1.303 | 3.998 | **<0.001** | **<0.001** |
| Action judgments: Low sound-high action vs. low sound-low action | 7.708 | 2.959 | 2.605 | *0.013* | *0.156* |
| **Mean RTs** | | | | | |
| Lexical decisions: Words vs. Pseudowords | -117.999 | 11.131 | -10.600 | **<0.001** | **<0.001** |
| Lexical decisions: High sound-high action vs. high sound-low action | 7.141 | 4.630 | 1.542 | 0.131 | 0.656 |
| Lexical decisions: High sound-high action vs. low sound-high action | 1.265 | 4.788 | 0.264 | 0.793 | 1.000 |
| Lexical decisions: High sound-high action vs. low sound-low action | -18.302 | 5.139 | -3.562 | **0.001** | **0.007** |
| Lexical decisions: High sound-low action vs. low sound-high action | -5.876 | 4.564 | -1.287 | 0.206 | 0.822 |
| Lexical decisions: High sound-low action vs. low sound-low action | -25.443 | 4.447 | -5.721 | **<0.001** | **<0.001** |
| Lexical decisions: Low sound-high action vs. low sound-low action | -19.567 | 4.762 | -4.109 | **<0.001** | **0.002** |
| Sound judgments: High sound-high action vs. high sound-low action | -138.208 | 12.006 | -11.511 | **<0.001** | **<0.001** |
| Sound judgments: High sound-high action vs. low sound-high action | -116.031 | 18.584 | -6.244 | **<0.001** | **<0.001** |
| Sound judgments: High sound-high action vs. low sound-low action | -60.796 | 16.696 | -3.641 | **0.001** | **0.006** |
| Sound judgments: High sound-low action vs. low sound-high action | 22.177 | 19.704 | 1.126 | 0.267 | 0.822 |
| Sound judgments: High sound-low action vs. low sound-low action | 77.412 | 18.638 | 4.154 | **<0.001** | **0.002** |
| Sound judgments: Low sound-high action vs. low sound-low action | 55.235 | 9.904 | 5.577 | **<0.001** | **<0.001** |
| Action judgments: High sound-high action vs. high sound-low action | -142.914 | 22.543 | -6.340 | **<0.001** | **<0.001** |
| Action judgments: High sound-high action vs. low sound-high action | -94.800 | 14.179 | -6.686 | **<0.001** | **<0.001** |
| Action judgments: High sound-high action vs. low sound-low action | -97.251 | 23.141 | -4.203 | **<0.001** | **0.001** |
| Action judgments: High sound-low action vs. low sound-high action | 48.114 | 21.937 | 2.193 | *0.034* | 0.206 |

| | | | | | |
|---|---|---|---|---|---|
| Action judgments: High sound-low action vs. low sound-low action | 45.662 | 11.470 | 3.981 | **<0.001** | **0.003** |
| Action judgments: Low sound-high action vs. low sound-low action | -2.452 | 20.606 | -0.119 | 0.906 | 1.000 |

**Table S7**. Study D (Martin et al. 2021): Repeated-measures ANOVA.

| **Effect** | **df** | **F** | **p** | **partial $\eta^2$** |
|---|---|---|---|---|
| **Error Rates** | | | | |
| TASK | 1,29 | 104.578 | **<0.001** | **0.783** |
| DIFFICULTY | 1,29 | 79.222 | **<0.001** | **0.732** |
| TASK x DIFFICULTY | 1,29 | 78.928 | **<0.001** | **0.731** |
| **Mean RTs** | | | | |
| TASK | 1,29 | 62.868 | **<0.001** | **0.684** |
| DIFFICULTY | 1,29 | 63.783 | **<0.001** | **0.687** |
| TASK x DIFFICULTY | 1,29 | 44.498 | **<0.001** | **0.605** |

**Table S8**. Study D (Martin et al. 2021): Post-hoc paired t-tests.

| **Comparison** | **Mean** | **SE** | **T** | **p (unc.)** | **p (corr.)** |
|---|---|---|---|---|---|
| **Error Rates** | | | | | |
| Semantic fluency vs. counting | 7.900 | 0.773 | 10.226 | **<0.001** | **<0.001** |
| Easy vs. difficult | -5.700 | 0.640 | -8.901 | **<0.001** | **<0.001** |
| Semantic fluency: easy vs. difficult | -11.067 | 1.200 | 9.226 | **<0.001** | **<0.001** |
| Counting: easy vs. difficult | -0.333 | 0.333 | 1.000 | 0.326 | 0.326 |
| **Mean RTs** | | | | | |
| Semantic fluency vs. counting | 111.902 | 14.113 | 7.929 | **<0.001** | **<0.001** |
| Easy vs. difficult | -56.620 | 7.090 | -7.986 | **<0.001** | **<0.001** |
| Semantic fluency: easy vs. difficult | -110.858 | 14.679 | 7.552 | **<0.001** | **<0.001** |
| Counting: easy vs. difficult | -2.383 | 4.157 | 0.573 | 0.571 | 0.571 |

**Table S9**. Study E (Turker et al. 2021): Repeated-measures ANOVA.

| **Effect** | **df** | **F** | **p** | **partial $\eta^2$** |
|---|---|---|---|---|
| **Error Rates** | | | | |
| WORD | 1,27 | 33.582 | **<0.001** | **0.554** |
| COMPLEXITY | 1,27 | 30.878 | **<0.001** | **0.533** |
| WORD x COMPLEXITY | 1,27 | 32.220 | **<0.001** | **0.544** |
| **Mean RTs** | | | | |
| WORD | 1,27 | 95.597 | **<0.001** | **0.780** |
| COMPLEXITY | 1,27 | 83.718 | **<0.001** | **0.756** |
| WORD x COMPLEXITY | 1,27 | 75.530 | **<0.001** | **0.737** |

**Table S10**. Study E (Turker et al. 2021): Post-hoc paired t-tests.

| **Comparison** | **Mean** | **SE** | **T** | **p (unc.)** | **p (corr.)** |
|---|---|---|---|---|---|
| **Error Rates** | | | | | |

| | | | | | |
|---|---|---|---|---|---|
| Words vs. pseudowords | -6.071 | 1.048 | -5.795 | **<0.001** | **<0.001** |
| Complex vs. simple | 4.071 | 0.733 | 5.557 | **<0.001** | **<0.001** |
| Complex vs. simple words | -0.143 | 0.340 | -0.420 | 0.678 | 0.678 |
| Complex vs. simple pseudowords | 8.286 | 1.435 | 5.772 | **<0.001** | **<0.001** |
| Words vs. pseudowords for complex vs. simple | -8.429 | 1.485 | -5.676 | **<0.001** | **<0.001** |
| **Mean RTs** | | | | | |
| Words vs. pseudowords | -201.207 | 20.579 | -9.777 | **<0.001** | **<0.001** |
| Complex vs. simple | 133.707 | 14.613 | 9.150 | **<0.001** | **<0.001** |
| Complex vs. simple words | 49.716 | 10.677 | 4.656 | **<0.001** | **<0.001** |
| Complex vs. simple pseudowords | 217.697 | 22.358 | 9.737 | **<0.001** | **<0.001** |
| Words vs. pseudowords for complex vs. simple | -167.981 | 19.329 | -8.691 | **<0.001** | **<0.001** |

### ROI analyses for each study

The following tables report the results of statistical analyses on the response profiles of the four AG-ROIs in each study. ROI responses were estimated as percent signal change vs. the resting baseline. For statistical inference, we first conducted one-sample t-tests to identify significant activation or deactivation, as compared to the resting baseline, for each experimental condition. Then, we tested for relative activity differences between different experimental conditions via repeated-measures ANOVA. Significant effects were resolved using step-down analyses and post-hoc paired t-tests. Bold font highlights significant effects ($p < 0.05$ Bonferroni-Holm corrected for multiple comparisons); italic fonts highlights trends ($p < 0.05$ uncorrected).

**Table S11.** Study A (Chapman and Hartwigsen 2021): One sample t-tests vs. the resting baseline.

| Task | Condition | Mean | SE | T | p (unc.) | p (corr.) |
|---|---|---|---|---|---|---|
| **Left PGa** | | | | | | |
| Taxonomic task | Taxonomic | -0.034 | 0.015 | -2.227 | *0.032* | *0.095* |
| Taxonomic task | Thematic | 0.002 | 0.013 | 0.133 | 0.895 | 1.000 |
| Taxonomic task | Unrelated | -0.076 | 0.014 | -5.635 | **<0.001** | **<0.001** |
| Thematic task | Taxonomic | -0.049 | 0.011 | -4.475 | **<0.001** | **<0.001** |
| Thematic task | Thematic | -0.001 | 0.013 | -0.069 | 0.945 | 1.000 |
| Thematic task | Unrelated | -0.097 | 0.012 | -7.984 | **<0.001** | **<0.001** |
| Scrambled task | Congruent | -0.174 | 0.015 | -11.325 | **<0.001** | **<0.001** |
| Scrambled task | Incongruent | -0.114 | 0.016 | -7.031 | **<0.001** | **<0.001** |
| **Left PGp** | | | | | | |
| Taxonomic task | Taxonomic | -0.003 | 0.019 | -0.153 | 0.879 | 1.000 |
| Taxonomic task | Thematic | 0.033 | 0.016 | 2.017 | 0.050 | 0.252 |

| | | | | | | |
|---|---|---|---|---|---|---|
| Taxonomic task | Unrelated | 0.006 | 0.014 | 0.402 | 0.690 | 1.000 |
| Thematic task | Taxonomic | -0.013 | 0.012 | -1.069 | 0.291 | 1.000 |
| Thematic task | Thematic | 0.052 | 0.016 | 3.304 | **0.002** | **0.012** |
| Thematic task | Unrelated | 0.007 | 0.013 | 0.504 | 0.617 | 1.000 |
| Scrambled task | Congruent | -0.112 | 0.019 | -5.846 | **0.000** | **0.000** |
| Scrambled task | Incongruent | -0.074 | 0.019 | -3.888 | **0.000** | **0.003** |
| **Right PGa** | | | | | | |
| Taxonomic task | Taxonomic | -0.047 | 0.015 | -3.156 | **0.003** | **0.012** |
| Taxonomic task | Thematic | -0.024 | 0.011 | -2.125 | *0.040* | *0.080* |
| Taxonomic task | Unrelated | -0.063 | 0.013 | -4.671 | **<0.001** | **<0.001** |
| Thematic task | Taxonomic | -0.039 | 0.012 | -3.138 | **0.003** | **0.012** |
| Thematic task | Thematic | -0.033 | 0.015 | -2.113 | *0.041* | *0.080* |
| Thematic task | Unrelated | -0.088 | 0.012 | -7.560 | **<0.001** | **<0.001** |
| Scrambled task | Congruent | -0.154 | 0.015 | -10.361 | **<0.001** | **<0.001** |
| Scrambled task | Incongruent | -0.103 | 0.013 | -8.003 | **<0.001** | **<0.001** |
| **Right PGp** | | | | | | |
| Taxonomic task | Taxonomic | -0.006 | 0.018 | -0.342 | 0.734 | 1.000 |
| Taxonomic task | Thematic | 0.014 | 0.015 | 0.951 | 0.347 | 1.000 |
| Taxonomic task | Unrelated | -0.025 | 0.016 | -1.488 | 0.145 | 0.723 |
| Thematic task | Taxonomic | -0.002 | 0.014 | -0.156 | 0.877 | 1.000 |
| Thematic task | Thematic | 0.015 | 0.017 | 0.888 | 0.380 | 1.000 |
| Thematic task | Unrelated | -0.039 | 0.014 | -2.786 | *0.008* | *0.057* |
| Scrambled task | Congruent | -0.079 | 0.019 | -4.224 | **<0.001** | **0.001** |
| Scrambled task | Incongruent | -0.046 | 0.018 | -2.535 | *0.015* | *0.091* |

**Table S12**. Study A (Chapman and Hartwigsen 2021): Repeated-measures ANOVA.

| **Effect** | **df** | **F** | **p** | **partial $\eta^2$** |
|---|---|---|---|---|
| **Full ANOVA** | | | | |
| ROI | 3,120 | 13.447 | **<0.001** | 0.252 |
| TASK | 1,40 | 0.239 | 0.628 | 0.006 |
| CONDITION | 2,80 | 32.462 | **<0.001** | 0.448 |
| ROI x TASK | 3,120 | 1.138 | 0.330 | 0.028 |
| ROI x CONDITION | 6,240 | 29.120 | **<0.001** | 0.421 |
| TASK x CONDITION | 2,80 | 0.599 | 0.540 | 0.015 |
| ROI x TASK x CONDITION | 6,240 | 4.310 | **0.001** | 0.097 |

| | | | | |
|---|---|---|---|---|
| **Left PGa** | | | | |
| TASK | 1,40 | 0.931 | 0.340 | 0.023 |
| CONDITION | 2,80 | 66.395 | **<0.001** | 0.624 |
| TASK x CONDITION | 2,80 | 0.486 | 0.597 | 0.012 |
| **Left PGp** | | | | |
| TASK | 1,40 | 0.077 | 0.783 | 0.002 |
| CONDITION | 2,80 | 20.109 | **<0.001** | 0.335 |
| TASK x CONDITION | 2,80 | 1.686 | 0.193 | 0.040 |
| **Right PGa** | | | | |
| TASK | 1,40 | 0.534 | 0.469 | 0.013 |
| CONDITION | 2,80 | 20.165 | **<0.001** | 0.335 |
| TASK x CONDITION | 2,80 | 2.055 | 0.135 | 0.049 |
| **Right PGp** | | | | |
| TASK | 1,40 | 0.069 | 0.794 | 0.002 |
| CONDITION | 2,80 | 21.422 | **<0.001** | 0.349 |
| TASK x CONDITION | 2,80 | 0.603 | 0.550 | 0.015 |

**Table S13**. Study A (Chapman and Hartwigsen 2021): Post-hoc paired t-tests.

| **Comparison** | **Mean** | **SE** | **T** | **p (unc.)** | **p (corr.)** |
|---|---|---|---|---|---|
| **Left PGa** | | | | | |
| Taxonomic vs. thematic conditions | -0.042 | 0.006 | -6.656 | **<0.001** | **<0.001** |
| Taxonomic vs. unrelated conditions | 0.045 | 0.008 | 5.593 | **<0.001** | **<0.001** |
| Thematic vs. unrelated conditions | 0.087 | 0.008 | 10.684 | **<0.001** | **<0.001** |
| Taxonomic vs. thematic task | 0.013 | 0.013 | 0.965 | 0.340 | 0.340 |
| Taxonomic vs. scrambled task | 0.107 | 0.016 | 6.858 | **<0.001** | **<0.001** |
| Thematic vs. scrambled task | 0.095 | 0.012 | 7.732 | **<0.001** | **<0.001** |
| Scrambled task: congruent vs. incongruent | -0.060 | 0.011 | -5.650 | **<0.001** | **<0.001** |
| **Left PGp** | | | | | |
| Taxonomic vs. thematic conditions | -0.050 | 0.008 | -6.678 | **<0.001** | **<0.001** |
| Taxonomic vs. unrelated conditions | -0.014 | 0.009 | -1.619 | 0.113 | 0.226 |
| Thematic vs. unrelated conditions | 0.036 | 0.008 | 4.438 | **<0.001** | **<0.001** |
| Taxonomic vs. thematic task | -0.003 | 0.012 | -0.278 | 0.783 | 0.783 |

| | | | | | |
|---|---|---|---|---|---|
| Taxonomic vs. scrambled task | 0.105 | 0.016 | 6.475 | **<0.001** | **<0.001** |
| Thematic vs. scrambled task | 0.108 | 0.014 | 7.622 | **<0.001** | **<0.001** |
| Scrambled task: congruent vs. incongruent | -0.038 | 0.010 | -3.847 | **<0.001** | **<0.001** |
| **Right PGa** | | | | | |
| Taxonomic vs. thematic conditions | -0.015 | 0.007 | -2.114 | *0.041* | *0.082* |
| Taxonomic vs. unrelated conditions | 0.033 | 0.008 | 4.192 | **<0.001** | **<0.001** |
| Thematic vs. unrelated conditions | 0.047 | 0.008 | 5.824 | **<0.001** | **<0.001** |
| Taxonomic vs. thematic task | 0.008 | 0.012 | 0.731 | 0.469 | 0.469 |
| Taxonomic vs. scrambled task | 0.084 | 0.015 | 5.409 | **<0.001** | **<0.001** |
| Thematic vs. scrambled task | 0.075 | 0.015 | 5.046 | **<0.001** | **<0.001** |
| Scrambled task: congruent vs. incongruent | -0.051 | 0.010 | -5.294 | **<0.001** | **<0.001** |
| **Right PGp** | | | | | |
| Taxonomic vs. thematic conditions | -0.019 | 0.006 | -3.009 | **0.005** | **0.012** |
| Taxonomic vs. unrelated conditions | 0.028 | 0.008 | 3.634 | **0.001** | **0.005** |
| Thematic vs. unrelated conditions | 0.046 | 0.007 | 6.210 | **<0.001** | **<0.001** |
| Taxonomic vs. thematic task | 0.003 | 0.012 | 0.263 | 0.794 | 0.794 |
| Taxonomic vs. scrambled task | 0.057 | 0.016 | 3.532 | **0.001** | **0.005** |
| Thematic vs. scrambled task | 0.054 | 0.014 | 3.953 | **<0.001** | **<0.001** |
| Scrambled task: congruent vs. incongruent | -0.033 | 0.011 | -3.057 | **0.004** | **0.012** |

**Table S14.** Study B (Graessner et al. 2021): One sample t-tests vs. the resting baseline.

| Task | Condition | Mean | SE | T | p (unc.) | p (corr.) |
|---|---|---|---|---|---|---|
| **Left PGa** | | | | | | |
| Implicit task | Meaningful phrases | -0.019 | 0.008 | -2.379 | *0.023* | *0.092* |
| Implicit task | Anomalous phrases | -0.019 | 0.007 | -2.631 | *0.013* | *0.065* |
| Implicit task | Pseudoword phrases | -0.037 | 0.009 | -3.950 | **<0.001** | **<0.001** |
| Implicit task | Single words | 0.006 | 0.007 | 0.822 | 0.417 | 0.834 |
| Explicit task | Meaningful phrases | 0.001 | 0.006 | 0.181 | 0.857 | 0.857 |
| Explicit task | Anomalous phrases | -0.020 | 0.007 | -2.920 | **0.006** | **0.036** |
| Explicit task | Pseudoword phrases | -0.033 | 0.006 | -5.945 | **<0.001** | **<0.001** |
| Explicit task | Single words | 0.010 | 0.006 | 1.755 | 0.089 | 0.267 |
| **Left PGp** | | | | | | |
| Implicit task | Meaningful phrases | -0.033 | 0.007 | -4.405 | **<0.001** | **<0.001** |
| Implicit task | Anomalous phrases | -0.037 | 0.009 | -4.257 | **<0.001** | **<0.001** |

| | | | | | | |
|---|---|---|---|---|---|---|
| Implicit task | Pseudoword phrases | -0.060 | 0.011 | -5.388 | **<0.001** | **<0.001** |
| Implicit task | Single words | -0.017 | 0.009 | -1.996 | 0.054 | 0.054 |
| Explicit task | Meaningful phrases | -0.018 | 0.007 | -2.746 | **0.010** | **0.020** |
| Explicit task | Anomalous phrases | -0.032 | 0.008 | -4.167 | **<0.001** | **<0.001** |
| Explicit task | Pseudoword phrases | -0.054 | 0.005 | -10.377 | **<0.001** | **<0.001** |
| Explicit task | Single words | -0.022 | 0.006 | -3.516 | **0.001** | **0.003** |
| **Right PGa** | | | | | | |
| Implicit task | Meaningful phrases | -0.003 | 0.007 | -0.461 | 0.648 | 0.894 |
| Implicit task | Anomalous phrases | -0.009 | 0.006 | -1.444 | 0.158 | 0.632 |
| Implicit task | Pseudoword phrases | -0.009 | 0.011 | -0.808 | 0.425 | 1.275 |
| Implicit task | Single words | 0.028 | 0.008 | 3.430 | **0.002** | **0.016** |
| Explicit task | Meaningful phrases | -0.006 | 0.007 | -0.771 | 0.447 | 1.275 |
| Explicit task | Anomalous phrases | -0.017 | 0.007 | -2.299 | *0.028* | *0.168* |
| Explicit task | Pseudoword phrases | -0.011 | 0.006 | -1.963 | 0.058 | 0.290 |
| Explicit task | Single words | 0.018 | 0.006 | 3.168 | **0.003** | **0.021** |
| **Right PGp** | | | | | | |
| Implicit task | Meaningful phrases | -0.035 | 0.006 | -5.355 | **<0.001** | **<0.001** |
| Implicit task | Anomalous phrases | -0.039 | 0.007 | -5.417 | **<0.001** | **<0.001** |
| Implicit task | Pseudoword phrases | -0.045 | 0.010 | -4.348 | **<0.001** | **<0.001** |
| Implicit task | Single words | -0.008 | 0.007 | -1.090 | 0.284 | 0.284 |
| Explicit task | Meaningful phrases | -0.028 | 0.009 | -3.214 | **0.003** | **0.009** |
| Explicit task | Anomalous phrases | -0.042 | 0.008 | -5.454 | **<0.001** | **<0.001** |
| Explicit task | Pseudoword phrases | -0.043 | 0.007 | -6.659 | **<0.001** | **<0.001** |
| Explicit task | Single words | -0.018 | 0.006 | -3.005 | **0.005** | **0.010** |

**Table S15.** Study B (Graessner et al. 2021): Repeated-measures ANOVA.

| **Effect** | **df** | **F** | **p** | **partial $\eta^2$** |
|---|---|---|---|---|
| **Full ANOVA** | | | | |
| ROI | 3,96 | 34.505 | **<0.001** | 0.519 |
| TASK | 1,32 | 0.054 | 0.818 | 0.002 |
| CONDITION | 3,96 | 30.543 | **<0.001** | 0.488 |
| ROI x TASK | 3,96 | 3.114 | **0.032** | 0.089 |
| ROI x CONDITION | 9,288 | 12.003 | **<0.001** | 0.273 |
| TASK x CONDITION | 3,96 | 1.714 | 0.183 | 0.051 |
| ROI x TASK x CONDITION | 9,288 | 2.020 | **0.048** | 0.059 |

| | | | | |
|---|---|---|---|---|
| **Left PGa** | | | | |
| TASK | 1,32 | 1.572 | 0.219 | 0.047 |
| CONDITION | 3,96 | 33.025 | **<0.001** | 0.508 |
| TASK x CONDITION | 3,96 | 3.472 | **0.019** | 0.098 |
| **Left PGp** | | | | |
| TASK | 1,32 | 0.598 | 0.445 | 0.018 |
| CONDITION | 3,96 | 27.031 | **<0.001** | 0.458 |
| TASK x CONDITION | 3,96 | 2.060 | 0.114 | 0.060 |
| **Right PGa** | | | | |
| TASK | 1,32 | 0.787 | 0.382 | 0.024 |
| CONDITION | 3,96 | 24.872 | **<0.001** | 0.437 |
| TASK x CONDITION | 3,96 | 0.511 | 0.629 | 0.016 |
| **Right PGp** | | | | |
| TASK | 1,32 | 0.028 | 0.868 | 0.001 |
| CONDITION | 3,96 | 20.098 | **<0.001** | 0.386 |
| TASK x CONDITION | 3,96 | 1.480 | 0.233 | 0.044 |

**Table S16.** Study B (Graessner et al. 2021): Post-hoc paired t-tests.

| Comparison | Mean | SE | T | p (unc.) | p (corr.) |
|---|---|---|---|---|---|
| **Left PGa** | | | | | |
| Implicit task: Meaningful vs. anomalous | 0.000 | 0.004 | 0.045 | 0.965 | 1.000 |
| Implicit task: Meaningful vs. pseudoword | 0.019 | 0.006 | 3.197 | **0.003** | **0.033** |
| Implicit task: Meaningful vs. single words | -0.025 | 0.005 | -4.925 | **<0.001** | **<0.001** |
| Implicit task: Anomalous vs. pseudoword | 0.019 | 0.007 | 2.784 | *0.009* | *0.081* |
| Implicit task: Anomalous vs. single words | -0.025 | 0.004 | -5.859 | **<0.001** | **<0.001** |
| Implicit task: Pseudoword vs. single words | -0.043 | 0.007 | -6.013 | **<0.001** | **<0.001** |
| Explicit task: Meaningful vs. anomalous | 0.021 | 0.006 | 3.723 | **0.001** | **0.012** |
| Explicit task: Meaningful vs. pseudoword | 0.034 | 0.006 | 5.583 | **<0.001** | **<0.001** |
| Explicit task: Meaningful vs. single words | -0.009 | 0.006 | -1.548 | 0.132 | 0.528 |
| Explicit task: Anomalous vs. pseudoword | 0.013 | 0.006 | 2.389 | *0.023* | *0.161* |
| Explicit task: Anomalous vs. single words | -0.030 | 0.005 | -5.926 | **<0.001** | **<0.001** |
| Explicit task: Pseudoword vs. single words | -0.043 | 0.006 | -7.548 | **<0.001** | **<0.001** |

| | | | | | |
|---|---|---|---|---|---|
| **Left PGp** | | | | | |
| Meaningful vs. anomalous phrases | 0.009 | 0.005 | 1.747 | 0.090 | 0.180 |
| Meaningful vs. pseudoword phrases | 0.031 | 0.004 | 8.553 | **<0.001** | **<0.001** |
| Meaningful phrases vs. single words | -0.006 | 0.005 | -1.413 | 0.167 | 0.180 |
| Anomalous vs. pseudoword phrases | 0.023 | 0.005 | 4.517 | **<0.001** | **<0.001** |
| Anomalous phrases vs. single words | -0.015 | 0.004 | -3.883 | **<0.001** | **<0.001** |
| Pseudowords phrases vs. single words | -0.038 | 0.005 | -8.073 | **<0.001** | **<0.001** |
| **Right PGa** | | | | | |
| Meaningful vs. anomalous phrases | 0.009 | 0.004 | 2.068 | *0.047* | *0.141* |
| Meaningful vs. pseudoword phrases | 0.005 | 0.003 | 1.581 | 0.124 | 0.248 |
| Meaningful phrases vs. single words | -0.027 | 0.005 | -5.276 | **<0.001** | **<0.001** |
| Anomalous vs. pseudoword phrases | -0.003 | 0.005 | -0.681 | 0.501 | 0.501 |
| Anomalous phrases vs. single words | -0.036 | 0.005 | -7.493 | **<0.001** | **<0.001** |
| Pseudowords phrases vs. single words | -0.033 | 0.005 | -6.023 | **<0.001** | **<0.001** |
| **Right PGp** | | | | | |
| Meaningful vs. anomalous phrases | 0.009 | 0.004 | 2.266 | *0.030* | *0.060* |
| Meaningful vs. pseudoword phrases | 0.013 | 0.004 | 3.673 | **0.001** | **0.004** |
| Meaningful phrases vs. single words | -0.019 | 0.005 | -3.747 | **0.001** | **0.004** |
| Anomalous vs. pseudoword phrases | 0.003 | 0.004 | 0.831 | 0.412 | 0.412 |
| Anomalous phrases vs. single words | -0.028 | 0.004 | -6.518 | **<0.001** | **<0.001** |
| Pseudowords phrases vs. single words | -0.032 | 0.005 | -5.886 | **<0.001** | **<0.001** |

**Table S17.** Study C (Kuhnke et al. 2020): One sample t-tests vs. the resting baseline.

| Task | Condition | Mean | SE | T | p (unc.) | p (corr.) |
|---|---|---|---|---|---|---|
| **Left PGa** | | | | | | |
| Lexical decision | High sound, high action | -0.023 | 0.012 | -1.855 | 0.071 | 0.712 |
| Lexical decision | High sound, low action | -0.011 | 0.012 | -0.891 | 0.378 | 1.000 |
| Lexical decision | Low sound, high action | 0.007 | 0.014 | 0.489 | 0.627 | 1.000 |
| Lexical decision | Low sound, low action | -0.027 | 0.015 | -1.853 | 0.072 | 0.712 |
| Lexical decision | Pseudowords | -0.080 | 0.013 | -6.301 | **<0.001** | **<0.001** |
| Sound judgment | High sound, high action | 0.027 | 0.014 | 1.974 | 0.056 | 0.611 |
| Sound judgment | High sound, low action | 0.020 | 0.016 | 1.251 | 0.218 | 1.000 |
| Sound judgment | Low sound, high action | -0.012 | 0.011 | -1.105 | 0.276 | 1.000 |
| Sound judgment | Low sound, low action | -0.019 | 0.015 | -1.320 | 0.194 | 1.000 |
| Action judgment | High sound, high action | 0.045 | 0.014 | 3.290 | **0.002** | **0.026** |

| | | | | | | |
|---|---|---|---|---|---|---|
| Action judgment | High sound, low action | 0.007 | 0.013 | 0.538 | 0.594 | 1.000 |
| Action judgment | Low sound, high action | 0.019 | 0.015 | 1.219 | 0.230 | 1.000 |
| Action judgment | Low sound, low action | -0.007 | 0.015 | -0.479 | 0.634 | 1.000 |
| **Left PGp** | | | | | | |
| Lexical decision | High sound, high action | -0.006 | 0.013 | -0.439 | 0.663 | 0.963 |
| Lexical decision | High sound, low action | 0.011 | 0.014 | 0.801 | 0.428 | 1.000 |
| Lexical decision | Low sound, high action | 0.015 | 0.015 | 0.995 | 0.326 | 1.000 |
| Lexical decision | Low sound, low action | -0.011 | 0.016 | -0.711 | 0.482 | 1.000 |
| Lexical decision | Pseudowords | -0.056 | 0.014 | -4.144 | **<0.001** | 0.002 |
| Sound judgment | High sound, high action | 0.038 | 0.014 | 2.692 | *0.010* | *0.104* |
| Sound judgment | High sound, low action | 0.042 | 0.014 | 2.930 | *0.006* | *0.062* |
| Sound judgment | Low sound, high action | 0.021 | 0.012 | 1.730 | 0.092 | 0.549 |
| Sound judgment | Low sound, low action | 0.024 | 0.015 | 1.606 | 0.116 | 0.581 |
| Action judgment | High sound, high action | 0.032 | 0.014 | 2.233 | *0.031* | *0.219* |
| Action judgment | High sound, low action | 0.032 | 0.014 | 2.320 | *0.026* | *0.215* |
| Action judgment | Low sound, high action | 0.036 | 0.015 | 2.352 | *0.024* | *0.215* |
| Action judgment | Low sound, low action | 0.041 | 0.013 | 3.069 | **0.004** | **0.047** |
| **Right PGa** | | | | | | |
| Lexical decision | High sound, high action | -0.009 | 0.013 | -0.683 | 0.499 | 0.998 |
| Lexical decision | High sound, low action | 0.004 | 0.012 | 0.374 | 0.710 | 0.998 |
| Lexical decision | Low sound, high action | 0.020 | 0.013 | 1.522 | 0.136 | 0.900 |
| Lexical decision | Low sound, low action | -0.016 | 0.012 | -1.313 | 0.197 | 0.984 |
| Lexical decision | Pseudowords | -0.042 | 0.011 | -3.868 | **<0.001** | **0.005** |
| Sound judgment | High sound, high action | -0.025 | 0.014 | -1.780 | 0.083 | 0.662 |
| Sound judgment | High sound, low action | -0.037 | 0.013 | -2.765 | *0.009* | *0.104* |
| Sound judgment | Low sound, high action | -0.019 | 0.012 | -1.553 | 0.129 | 0.900 |
| Sound judgment | Low sound, low action | -0.017 | 0.014 | -1.283 | 0.207 | 0.984 |
| Action judgment | High sound, high action | -0.015 | 0.013 | -1.180 | 0.245 | 0.828 |
| Action judgment | High sound, low action | -0.026 | 0.014 | -1.874 | 0.068 | 0.631 |
| Action judgment | Low sound, high action | -0.035 | 0.014 | -2.455 | *0.019* | *0.205* |
| Action judgment | Low sound, low action | -0.025 | 0.013 | -1.913 | 0.063 | 0.631 |
| **Right PGp** | | | | | | |
| Lexical decision | High sound, high action | -0.031 | 0.011 | -2.824 | *0.007* | *0.089* |
| Lexical decision | High sound, low action | -0.006 | 0.011 | -0.562 | 0.577 | 1.000 |

| | | | | | | |
|---|---|---|---|---|---|---|
| Lexical decision | Low sound, high action | -0.003 | 0.012 | -0.219 | 0.828 | 1.000 |
| Lexical decision | Low sound, low action | -0.028 | 0.012 | -2.323 | *0.026* | *0.230* |
| Lexical decision | Pseudowords | -0.053 | 0.011 | -4.943 | **<0.001** | **<0.001** |
| Sound judgment | High sound, high action | -0.020 | 0.012 | -1.646 | 0.108 | 0.862 |
| Sound judgment | High sound, low action | -0.030 | 0.012 | -2.573 | *0.014* | *0.140* |
| Sound judgment | Low sound, high action | -0.017 | 0.012 | -1.462 | 0.152 | 1.000 |
| Sound judgment | Low sound, low action | -0.012 | 0.013 | -0.907 | 0.370 | 1.000 |
| Action judgment | High sound, high action | -0.014 | 0.012 | -1.145 | 0.259 | 1.000 |
| Action judgment | High sound, low action | -0.018 | 0.012 | -1.479 | 0.147 | 1.000 |
| Action judgment | Low sound, high action | -0.034 | 0.013 | -2.643 | *0.012* | *0.130* |
| Action judgment | Low sound, low action | -0.015 | 0.012 | -1.272 | 0.211 | 1.000 |

**Table S18.** Study C (Kuhnke et al. 2020): Repeated-measures ANOVA.

| **Effect** | **df** | **F** | **p** | **partial $\eta^2$** |
|---|---|---|---|---|
| **Full ANOVA** | | | | |
| ROI | 3,117 | 10.429 | **<0.001** | 0.211 |
| TASK | 2,78 | 0.327 | 0.705 | 0.008 |
| SOUND | 1,39 | 1.353 | 0.252 | 0.034 |
| ACTION | 1,39 | 1.515 | 0.226 | 0.037 |
| ROI x TASK | 6,234 | 12.386 | **<0.001** | 0.241 |
| ROI x SOUND | 3,117 | 12.536 | **<0.001** | 0.243 |
| TASK x SOUND | 2,78 | 1.040 | 0.358 | 0.026 |
| ROI x TASK x SOUND | 6,234 | 9.546 | **<0.001** | 0.197 |
| ROI x ACTION | 3,117 | 6.650 | **<0.001** | 0.146 |
| TASK x ACTION | 2,78 | 0.102 | 0.900 | 0.003 |
| ROI x TASK x ACTION | 6,234 | 2.580 | **0.028** | 0.062 |
| SOUND x ACTION | 1,39 | 1.577 | 0.217 | 0.039 |
| ROI x SOUND x ACTION | 3,117 | 0.716 | 0.542 | 0.018 |
| TASK x SOUND x ACTION | 2,78 | 4.345 | **0.019** | 0.100 |
| ROI x TASK x SOUND x ACTION | 6,234 | 0.577 | 0.738 | 0.015 |
| **Left PGa** | | | | |
| TASK | 2,78 | 3.058 | 0.055 | 0.073 |
| SOUND | 1,39 | 12.946 | **0.001** | 0.249 |
| ACTION | 1,39 | 9.376 | **0.004** | 0.194 |
| TASK x SOUND | 2,78 | 7.104 | **0.001** | 0.154 |
| TASK x ACTION | 2,78 | 2.253 | 0.112 | 0.055 |

| | | | | |
|---|---|---|---|---|
| SOUND x ACTION | 2,78 | 3.058 | 0.055 | 0.073 |
| **Left PGp** | | | | |
| TASK | 2,78 | 4.365 | **0.021** | 0.101 |
| SOUND | 1,39 | 0.735 | 0.397 | 0.018 |
| ACTION | 1,39 | 0.007 | 0.933 | 0.000 |
| TASK x SOUND | 2,78 | 2.324 | 0.105 | 0.056 |
| TASK x ACTION | 2,78 | 0.347 | 0.705 | 0.009 |
| SOUND x ACTION | 2,78 | 4.365 | **0.021** | 0.101 |
| **Right PGa** | | | | |
| TASK | 2,78 | 2.809 | 0.071 | 0.067 |
| SOUND | 1,39 | 0.411 | 0.525 | 0.010 |
| ACTION | 1,39 | 1.487 | 0.230 | 0.037 |
| TASK x SOUND | 2,78 | 2.043 | 0.137 | 0.050 |
| TASK x ACTION | 2,78 | 0.431 | 0.652 | 0.011 |
| SOUND x ACTION | 2,78 | 2.809 | 0.071 | 0.067 |
| **Right PGp** | | | | |
| TASK | 2,78 | 0.054 | 0.947 | 0.001 |
| SOUND | 1,39 | 0.318 | 0.576 | 0.008 |
| ACTION | 1,39 | 0.140 | 0.710 | 0.004 |
| TASK x SOUND | 2,78 | 2.038 | 0.137 | 0.050 |
| TASK x ACTION | 2,78 | 0.381 | 0.666 | 0.010 |
| SOUND x ACTION | 2,78 | 0.054 | 0.947 | 0.001 |

**Table S19.** Study C (Kuhnke et al. 2020): Post-hoc paired t-tests.

| Comparison | Mean | SE | T | p (unc.) | p (corr.) |
|---|---|---|---|---|---|
| **Left PGa** | | | | | |
| Lexical decision: Words vs. Pseudowords | 0.067 | 0.008 | 8.021 | **<0.001** | **<0.001** |
| Lexical decision: High- vs. low-sound words | -0.006 | 0.009 | -0.745 | 0.461 | 0.922 |
| Lexical decision: High- vs. low-action words | 0.011 | 0.008 | 1.393 | 0.171 | 0.513 |
| Sound judgment: High- vs. low-sound words | 0.039 | 0.009 | 4.424 | **<0.001** | **<0.001** |
| Sound judgment: High- vs. low-action words | 0.007 | 0.009 | 0.732 | 0.469 | 0.922 |
| Sound judgment: High- vs. low-action words | 0.032 | 0.010 | 3.161 | **0.003** | **0.015** |
| Sound judgment: High- vs. low-sound words | 0.021 | 0.008 | 2.446 | *0.019* | *0.076* |

| | | | | | |
|---|---|---|---|---|---|
| **Left PGp** | | | | | |
| Lexical decision: Words vs. Pseudowords | 0.058 | 0.008 | 7.210 | **<0.001** | **<0.001** |
| Lexical decisions vs. Sound judgments | -0.029 | 0.012 | -2.376 | *0.023* | *0.069* |
| Lexical decisions vs. Action judgments | -0.033 | 0.014 | -2.289 | *0.028* | *0.069* |
| Sound judgments vs. Action judgments | -0.004 | 0.009 | -0.428 | 0.671 | 0.671 |
| **Right PGa** | | | | | |
| Lexical decision: Words vs. Pseudowords | 0.041 | 0.009 | 4.635 | **<0.001** | **<0.001** |
| Lexical decision: High- vs. low-sound words | -0.004 | 0.007 | -0.571 | 0.571 | 1.000 |
| Lexical decision: High- vs. low-action words | 0.011 | 0.008 | 1.440 | 0.158 | 0.858 |
| Sound judgment: High- vs. low-sound words | -0.013 | 0.009 | -1.496 | 0.143 | 0.858 |
| Sound judgment: High- vs. low-action words | 0.005 | 0.008 | 0.625 | 0.536 | 1.000 |
| Sound judgment: High- vs. low-action words | 0.000 | 0.009 | 0.045 | 0.964 | 1.000 |
| Sound judgment: High- vs. low-sound words | 0.009 | 0.007 | 1.338 | 0.189 | 0.790 |
| **Right PGp** | | | | | |
| Lexical decision: Words vs. Pseudowords | 0.036 | 0.007 | 5.442 | **<0.001** | **<0.001** |
| Lexical decision: High- vs. low-sound words | -0.004 | 0.007 | -0.539 | 0.593 | 1.000 |
| Lexical decision: High- vs. low-action words | 0.000 | 0.008 | -0.010 | 0.992 | 1.606 |
| Sound judgment: High- vs. low-sound words | -0.011 | 0.007 | -1.557 | 0.128 | 0.768 |
| Sound judgment: High- vs. low-action words | 0.002 | 0.008 | 0.251 | 0.803 | 1.000 |
| Sound judgment: High- vs. low-action words | -0.007 | 0.008 | -0.911 | 0.368 | 1.000 |
| Sound judgment: High- vs. low-sound words | 0.008 | 0.006 | 1.374 | 0.177 | 0.885 |

**Table S20.** Study D (Martin et al. 2021): One sample t-tests vs. the resting baseline.

| Task | Condition | Mean | SE | T | p (unc.) | p (corr.) |
|---|---|---|---|---|---|---|
| **Left PGa** | | | | | | |
| Semantic fluency | Difficult | -0.367 | 0.036 | -10.077 | **<0.001** | **<0.001** |
| Semantic fluency | Easy | -0.285 | 0.033 | -8.716 | **<0.001** | **<0.001** |
| Counting | Difficult | -0.233 | 0.038 | -6.182 | **<0.001** | **<0.001** |
| Counting | Easy | -0.188 | 0.033 | -5.650 | **<0.001** | **<0.001** |
| **Left PGp** | | | | | | |
| Semantic fluency | Difficult | -0.265 | 0.031 | -8.416 | **<0.001** | **<0.001** |
| Semantic fluency | Easy | -0.196 | 0.031 | -6.390 | **<0.001** | **<0.001** |
| Counting | Difficult | -0.202 | 0.030 | -6.841 | **<0.001** | **<0.001** |
| Counting | Easy | -0.139 | 0.023 | -5.969 | **<0.001** | **<0.001** |

| | | | | | | |
|---|---|---|---|---|---|---|
| **Right PGa** | | | | | | |
| Semantic fluency | Difficult | -0.428 | 0.053 | -8.072 | **<0.001** | **<0.001** |
| Semantic fluency | Easy | -0.298 | 0.045 | -6.563 | **<0.001** | **<0.001** |
| Counting | Difficult | -0.155 | 0.028 | -5.621 | **<0.001** | **<0.001** |
| Counting | Easy | -0.160 | 0.034 | -4.753 | **<0.001** | **<0.001** |
| **Right PGp** | | | | | | |
| Semantic fluency | Difficult | -0.494 | 0.052 | -9.465 | **<0.001** | **<0.001** |
| Semantic fluency | Easy | -0.311 | 0.040 | -7.703 | **<0.001** | **<0.001** |
| Counting | Difficult | -0.188 | 0.024 | -7.840 | **<0.001** | **<0.001** |
| Counting | Easy | -0.162 | 0.026 | -6.345 | **<0.001** | **<0.001** |

**Table S21.** Study D (Martin et al. 2021): Repeated-measures ANOVA.

| **Effect** | **df** | **F** | **p** | **partial $\eta^2$** |
|---|---|---|---|---|
| **Full ANOVA** | | | | |
| ROI | 3,87 | 8.243 | **<0.001** | 0.221 |
| TASK | 1,29 | 23.292 | **<0.001** | 0.445 |
| DIFFICULTY | 1,29 | 8.691 | **0.006** | 0.231 |
| ROI x TASK | 3,87 | 9.901 | **<0.001** | 0.255 |
| ROI x DIFFICULTY | 3,87 | 3.944 | **0.013** | 0.120 |
| TASK x DIFFICULTY | 1,29 | 6.571 | **0.016** | 0.185 |
| ROI x TASK x DIFFICULTY | 3,87 | 5.446 | **0.008** | 0.158 |
| **Left PGa** | | | | |
| TASK | 1,29 | 8.334 | **0.007** | 0.223 |
| DIFFICULTY | 1,29 | 7.346 | **0.011** | 0.202 |
| TASK x DIFFICULTY | 1,29 | 0.733 | 0.399 | 0.025 |
| **Left PGp** | | | | |
| TASK | 1,29 | 4.268 | **0.048** | 0.128 |
| DIFFICULTY | 1,29 | 7.958 | **0.009** | 0.215 |
| TASK x DIFFICULTY | 1,29 | 0.029 | 0.865 | 0.001 |
| **Right PGa** | | | | |
| TASK | 1,29 | 27.069 | **<0.001** | 0.483 |
| DIFFICULTY | 1,29 | 4.893 | **0.035** | 0.144 |

| | | | | |
|---|---|---|---|---|
| TASK x DIFFICULTY | 1,29 | 6.410 | **0.017** | 0.181 |
| **Right PGp** | | | | |
| TASK | 1,29 | 28.245 | **<0.001** | 0.493 |
| DIFFICULTY | 1,29 | 11.589 | **0.002** | 0.286 |
| TASK x DIFFICULTY | 1,29 | 20.217 | **<0.001** | 0.411 |

**Table S22.** Study D (Martin et al. 2021): Post-hoc paired t-tests.

| Comparison | Mean | SE | T | p (unc.) | p (corr.) |
|---|---|---|---|---|---|
| **Left PGa** | | | | | |
| Semantic fluency vs. counting | -0.115 | 0.040 | -2.887 | **0.007** | **0.014** |
| Easy vs. difficult | 0.064 | 0.024 | 2.710 | **0.011** | **0.011** |
| **Left PGp** | | | | | |
| Semantic fluency vs. counting | -0.060 | 0.029 | -2.066 | **0.048** | **0.048** |
| Easy vs. difficult | 0.066 | 0.023 | 2.821 | **0.009** | **0.018** |
| **Right PGa** | | | | | |
| Semantic fluency vs. counting | -0.206 | 0.040 | -5.203 | **<0.001** | **<0.001** |
| Easy vs. difficult | 0.063 | 0.028 | 2.212 | *0.035* | *0.070* |
| Semantic fluency: easy vs. difficult | 0.130 | 0.046 | 2.851 | **0.008** | **0.024** |
| Counting: easy vs. difficult | -0.004 | 0.031 | -0.132 | 0.896 | 0.896 |
| **Right PGp** | | | | | |
| Semantic fluency vs. counting | -0.228 | 0.043 | -5.315 | **<0.001** | **<0.001** |
| Easy vs. difficult | 0.105 | 0.031 | 3.404 | **0.002** | **0.004** |
| Semantic fluency: easy vs. difficult | 0.183 | 0.039 | 4.668 | **<0.001** | **<0.001** |
| Counting: easy vs. difficult | 0.027 | 0.031 | 0.857 | 0.399 | 0.399 |

**Table S23.** Study E (Turker et al. 2021): One sample t-tests vs. the resting baseline.

| Condition | Mean | SE | T | p (unc.) | p (corr.) |
|---|---|---|---|---|---|
| **Left PGa** | | | | | |
| Complex words | -0.025 | 0.021 | -1.223 | 0.232 | 0.232 |
| Simple words | 0.029 | 0.017 | 1.705 | 0.100 | 0.199 |
| Complex pseudowords | -0.235 | 0.028 | -8.525 | **<0.001** | **<0.001** |
| Simple pseudowords | -0.075 | 0.020 | -3.670 | **0.001** | **0.003** |

| | | | | | |
|---|---|---|---|---|---|
| **Left PGp** | | | | | |
| Complex words | -0.019 | 0.019 | -0.960 | 0.345 | 0.460 |
| Simple words | 0.016 | 0.013 | 1.229 | 0.230 | 0.460 |
| Complex pseudowords | -0.159 | 0.027 | -5.826 | **<0.001** | **<0.001** |
| 16Simple pseudowords | -0.080 | 0.019 | -4.227 | **<0.001** | **0.001** |
| **Right PGa** | | | | | |
| Complex words | -0.044 | 0.022 | -2.002 | 0.055 | 0.111 |
| Simple words | -0.009 | 0.016 | -0.561 | 0.579 | 0.579 |
| Complex pseudowords | -0.250 | 0.028 | -8.826 | **<0.001** | **<0.001** |
| Simple pseudowords | -0.097 | 0.027 | -3.539 | **0.001** | **0.004** |
| **Right PGp** | | | | | |
| Complex words | -0.054 | 0.018 | -3.053 | **0.005** | **0.010** |
| Simple words | -0.019 | 0.015 | -1.236 | 0.227 | 0.227 |
| Complex pseudowords | -0.198 | 0.027 | -7.286 | **<0.001** | **<0.001** |
| Simple pseudowords | -0.096 | 0.021 | -4.670 | **<0.001** | **<0.001** |

**Table S24.** Study E (Turker et al. 2021): Repeated-measures ANOVA.

| Effect | df | F | p | partial $\eta^2$ |
|---|---|---|---|---|
| **Full ANOVA** | | | | |
| ROI | 3,81 | 2.892 | **0.049** | 0.097 |
| WORD | 1,27 | 132.208 | **<0.001** | 0.830 |
| ROI x WORD | 3,81 | 4.087 | **0.018** | 0.131 |
| ROI x COMPLEXITY | 3,81 | 10.073 | **<0.001** | 0.272 |
| WORD x COMPLEXITY | 1,27 | 10.034 | **0.004** | 0.271 |
| ROI x WORD x COMPLEXITY | 3,81 | 6.926 | **0.001** | 0.204 |
| **Left PGa** | | | | |
| WORD | 1,27 | 95.790 | **<0.001** | 0.780 |
| COMPLEXITY | 1,27 | 58.472 | **<0.001** | 0.684 |
| WORD x COMPLEXITY | 1,27 | 14.446 | **0.001** | 0.349 |
| **Left PGp** | | | | |
| WORD | 1,27 | 110.783 | **<0.001** | 0.804 |
| COMPLEXITY | 1,27 | 16.148 | **<0.001** | 0.374 |
| WORD x COMPLEXITY | 1,27 | 3.484 | 0.073 | 0.114 |

| | | | | |
|---|---|---|---|---|
| **Right PGa** | | | | |
| WORD | 1,27 | 72.858 | **<0.001** | 0.730 |
| COMPLEXITY | 1,27 | 52.889 | **<0.001** | 0.662 |
| WORD x COMPLEXITY | 1,27 | 14.596 | **0.001** | 0.351 |
| **Right PGp** | | | | |
| WORD | 1,27 | 57.379 | **<0.001** | 0.680 |
| COMPLEXITY | 1,27 | 23.459 | **<0.001** | 0.465 |
| WORD x COMPLEXITY | 1,27 | 4.441 | **0.045** | 0.141 |

**Table S25.** Study E (Turker et al. 2021): Post-hoc paired t-tests.

| Comparison | Mean | SE | T | p (unc.) | p (corr.) |
|---|---|---|---|---|---|
| **Left PGa** | | | | | |
| Words vs. pseudowords | 0.157 | 0.016 | 9.787 | **<0.001** | **<0.001** |
| Complex vs. simple | -0.108 | 0.014 | -7.647 | **<0.001** | **<0.001** |
| Complex vs. simple words | -0.054 | 0.016 | -3.499 | **0.002** | **0.002** |
| Complex vs. simple pseudowords | -0.161 | 0.023 | -6.876 | **<0.001** | **<0.001** |
| Words vs. pseudowords for complex vs. simple | 0.106 | 0.028 | 3.801 | **0.001** | **0.002** |
| **Left PGp** | | | | | |
| Words vs. pseudowords | 0.118 | 0.011 | 10.525 | **<0.001** | **<0.001** |
| Complex vs. simple | -0.057 | 0.014 | -4.018 | **<0.001** | **<0.001** |
| Complex vs. simple words | -0.035 | 0.014 | -2.467 | **0.020** | **0.040** |
| Complex vs. simple pseudowords | -0.079 | 0.022 | -3.583 | **0.001** | **0.003** |
| Words vs. pseudowords for complex vs. simple | 0.045 | 0.024 | 1.866 | 0.073 | 0.073 |
| **Right PGa** | | | | | |
| Words vs. pseudowords | 0.147 | 0.017 | 8.536 | **<0.001** | **<0.001** |
| Complex vs. simple | -0.094 | 0.013 | -7.272 | **<0.001** | **<0.001** |
| Complex vs. simple words | -0.035 | 0.020 | -1.791 | 0.085 | 0.085 |
| Complex vs. simple pseudowords | -0.153 | 0.021 | -7.371 | **<0.001** | **<0.001** |
| Words vs. pseudowords for complex vs. simple | 0.118 | 0.031 | 3.820 | **0.001** | **0.002** |
| **Right PGp** | | | | | |
| Words vs. pseudowords | 0.111 | 0.015 | 7.575 | **<0.001** | **<0.001** |
| Complex vs. simple | -0.068 | 0.014 | -4.843 | **<0.001** | **<0.001** |

| | | | | | |
|---|---|---|---|---|---|
| Complex vs. simple words | -0.035 | 0.017 | -2.067 | *0.048* | *0.090* |
| Complex vs. simple pseudowords | -0.102 | 0.025 | -4.083 | **<0.001** | **<0.001** |
| Words vs. pseudowords for complex vs. simple | 0.067 | 0.032 | 2.107 | *0.045* | *0.090* |

### Linear-mixed-model analysis across all studies

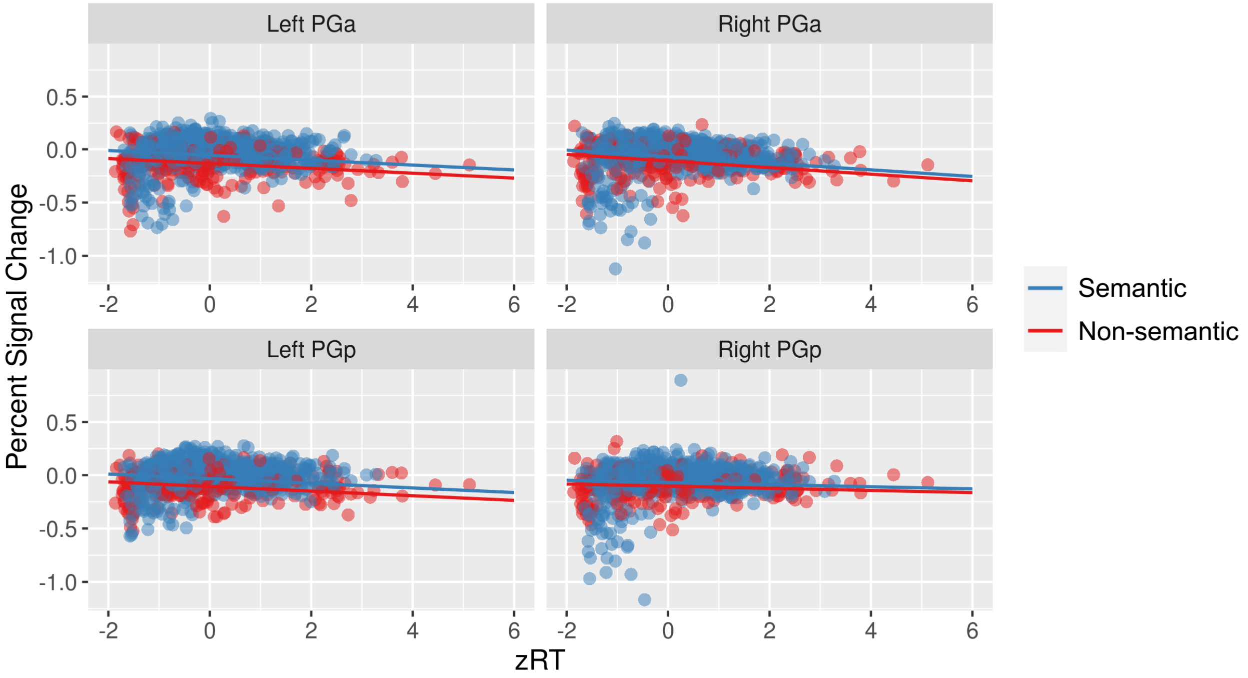

**Figure S1. Results of the linear-mixed-model analysis.** Combined effects of semantics and zRT on predicted percent signal change in each ROI, including raw data points.

**Table S26.** ANOVA table of the next best-fitting model that included the zRT x Semantics interaction.

| Factor | SS | MSE | DF | Den. DF | F | p |
|---|---|---|---|---|---|---|
| zRT | 0.32 | 0.32 | 1 | 4876.6 | 39.63 | **< 0.001** |
| Sem | 0.09 | 0.09 | 1 | 27.9 | 10.96 | **0.003** |
| ROI | 0.35 | 0.12 | 3 | 5187.4 | 14.67 | **< 0.001** |
| zRT x ROI | 0.29 | 0.10 | 3 | 5187.4 | 12.18 | **< 0.001** |
| Sem x ROI | 0.33 | 0.11 | 3 | 5187.4 | 13.68 | **< 0.001** |
| zRT x Sem | 0.01 | 0.01 | 1 | 5006.1 | 1.69 | 0.194 |

SS = sum of squares; MSE = mean squared error; DF = degrees of freedom; Den = denominator; Sem = effect of semantics.

**Table S27.** Fixed effects of mixed model with PGa vs. PGp and left vs. right contrasts.

| | Estimate | SE | df | t | p |
|---|---|---|---|---|---|
| Intercept | -0.085 | 0.031 | 11.6 | -2.74 | **0.018** |
| zRT | -0.021 | 0.003 | 4812.7 | -6.74 | **<0.001** |
| Sem vs. Non-sem | 0.056 | 0.017 | 27.7 | 3.39 | **0.002** |
| ROI: PGa vs. PGp | -0.031 | 0.006 | 5187.3 | -5.22 | **<0.001** |
| ROI: Left vs. Right | 0.011 | 0.006 | 5187.3 | 1.84 | 0.066 |
| zRT x PGa vs. PGp | -0.022 | 0.005 | 5187.3 | -4.44 | **<0.001** |
| zRT x Left vs. Right | -0.004 | 0.005 | 5187.3 | -0.72 | 0.470 |
| Sem vs. Non-sem x PGa vs. PGp | 0.009 | 0.012 | 5187.3 | 0.74 | 0.460 |
| Sem vs. Non-sem x Left vs. Right | 0.075 | 0.012 | 5187.3 | 6.33 | **<0.001** |

**Table S28.** Fixed effects of left PGa model.

| | **Estimate** | **SE** | **df** | **t** | **p** |
|---|---|---|---|---|---|
| Intercept | -0.13 | 0.033 | 16.4 | -3.94 | **0.001** |
| zRT | -0.01 | 0.006 | 1071.1 | -2.43 | **0.015** |
| Sem vs. Non-sem | 0.07 | 0.021 | 29.1 | 3.53 | **0.001** |

**Table S29.** Fixed effects of left PGp model.

| | **Estimate** | **SE** | **df** | **t** | **p** |
|---|---|---|---|---|---|
| Intercept | -0.09 | 0.028 | 13.9 | -3.32 | **0.005** |
| zRT | -0.02 | 0.005 | 1108.9 | -4.75 | **<0.001** |
| Sem vs. Non-sem | 0.06 | 0.013 | 27.5 | 4.51 | **<0.001** |

**Table S30.** Fixed effects of right PGa model.

| | **Estimate** | **SE** | **df** | **t** | **p** |
|---|---|---|---|---|---|
| Intercept | -0.12 | 0.036 | 14.6 | -3.23 | **0.006** |
| zRT | -0.03 | 0.006 | 1189.2 | -4.60 | **<0.001** |
| Sem vs. Non-sem | 0.05 | 0.019 | 27.3 | 2.68 | **0.012** |

**Table S31.** Fixed effects of right PGp model.

| | **Estimate** | **SE** | **df** | **t** | **p** |
|---|---|---|---|---|---|
| Intercept | -0.11 | 0.040 | 13.6 | -2.88 | **0.013** |
| zRT | -0.02 | 0.006 | 1230.3 | -3.71 | **<0.001** |
| Sem vs. Non-sem | 0.04 | 0.018 | 26.8 | 2.45 | **0.021** |

**Separate linear-mixed-model analyses for semantic and non-semantic conditions**

To test whether RT effects in the AG are indeed domain-general and emerge in both semantic and non-semantic conditions, we ran two additional linear-mixed-model analyses for semantic and non-semantic conditions separately.

The results show that the optimal model of AG responses included main effects of zRT and ROI, as well as interactions of zRT x ROI for both semantic conditions (Tables S32-S33) and non-semantic conditions (Tables S34-S35). These findings support our conclusion that effects of RT on AG activity are domain-general and independent of semantics.

*Semantic conditions only*

**Table S32**. AIC comparisons to determine the optimal model predicting AG activity for semantic conditions only.

| Comparison | Model | ΔAIC | df | |
|---|---|---|---|---|
| **(A) Random effects** | **PSC ~ 1 + (1 \| participant) + (1 \| task/condition)** | **0** | **5** | * |
| | PSC ~ 1 + (1 \| participant) + (1 \| condition) | 18.4 | 4 | |
| | PSC ~ 1 + (1 \| participant) + (1 \| task) | 140.4 | 4 | |
| | PSC ~ 1 + (1 \| participant) | 985.9 | 3 | |
| **(B) zRT & zER** | **PSC ~ zRT + (1 \| participant) + (1 \| task/condition)** | **0** | **6** | * |
| | PSC ~ zER + (1 \| participant) + (1 \| task/condition) | 28.9 | 6 | |
| | PSC ~ 1 + (1 \| participant) + (1 \| task/condition) | 32.0 | 5 | |
| **(C) Interactions** | **PSC ~ zRT + ROI + zRT:ROI + (1 \| participant) + (1 \| task/condition)** | **0** | **12** | * |
| | PSC ~ zRT + ROI + (1 \| participant) + (1 \| task/condition) | 20.3 | 9 | |
| | PSC ~ zRT + (1 \| participant) + (1 \| task/condition) | 140.3 | 6 | |

* = winning model; PSC = percent signal change; zRT = standardized RT; zER = standardized error rate; AIC = Akaike Information Criterion.

**Table S33.** ANOVA table for the optimal model of semantic conditions.

| Factor | SS | MSE | DF | Den. DF | F | p |
|---|---|---|---|---|---|---|
| zRT | 0.22 | 0.22 | 1 | 3821.3 | 30.33 | **< 0.001** |
| ROI | 0.91 | 0.30 | 3 | 3983.8 | 42.70 | **< 0.001** |
| zRT x ROI | 0.19 | 0.06 | 3 | 3983.8 | 8.80 | **< 0.001** |

SS = sum of squares; MSE = mean squared error; DF = degrees of freedom; Den = denominator.

*Non-semantic conditions only*

**Table S34**. AIC comparisons to determine the optimal model predicting AG activity for non-semantic conditions only.

| Comparison | Model | ΔAIC | df | |
|---|---|---|---|---|
| **(A) Random effects** | **PSC ~ 1 + (1 \| participant) + (1 \| condition)** | **0** | **4** | * |
| | PSC ~ 1 + (1 \| participant) + (1 \| task/condition) | 6.6 | 5 | |
| | PSC ~ 1 + (1 \| participant) + (1 \| task) | 106.8 | 4 | |
| | PSC ~ 1 + (1 \| participant) | 252.6 | 3 | |
| **(B) zRT & zER** | **PSC ~ zRT + (1 \| participant) + (1 \| condition)** | **0** | **5** | * |
| | PSC ~ zER + (1 \| participant) + (1 \| task/condition) | 11.1 | 5 | |
| | PSC ~ 1 + (1 \| participant) + (1 \| task/condition) | 14.7 | 4 | |
| **(C) Interactions** | **PSC ~ zRT + ROI + zRT:ROI + (1 \| participant) + (1 \| condition)** | **0** | **11** | * |
| | PSC ~ zRT + ROI + (1 \| participant) + (1 \| condition) | 7.6 | 8 | |
| | PSC ~ zRT + (1 \| participant) + (1 \| condition) | 23.3 | 5 | |

* = winning model; PSC = percent signal change; zRT = standardized RT; zER = standardized error rate; AIC = Akaike Information Criterion.

**Table S35.** ANOVA table for the optimal model of non-semantic conditions.

| Factor | SS | MSE | DF | Den. DF | F | p |
|---|---|---|---|---|---|---|
| zRT | 0.10 | 0.10 | 1 | 763.7 | 12.14 | **< 0.001** |
| ROI | 0.17 | 0.06 | 3 | 1084.8 | 6.86 | **< 0.001** |
| zRT x ROI | 0.11 | 0.04 | 3 | 1084.8 | 4.55 | **0.004** |

SS = sum of squares; MSE = mean squared error; DF = degrees of freedom; Den = denominator.

### Effect of semantics on RT

To investigate the relationship between the semantics and RT variables in our model, we ran a supplementary linear-mixed-model analysis that aimed to explain zRT by semantics and random effects as in our original model (i.e. subject and condition nested within task):

$$zRT \sim Sem + (1 \mid participant) + (1 \mid task/condition)$$

The results show that the effect of semantics on zRT is significantly different from zero (Table S36), but very small, explaining only 1.1% of the variance in RT (Table S37, $R^2$ marg.). Note that the entire model explains 89.1% of the variance in RT (Table S37, $R^2$ cond.).

**Table S36.** ANOVA table of the linear-mixed-effects model explaining zRT by semantics.

| Factor | SS | MSE | DF | Den. DF | F | p |
|---|---|---|---|---|---|---|
| Sem | 0.81 | 0.81 | 1 | 26.16 | 5.69 | **0.025** |

SS = sum of squares; MSE = mean squared error; DF = degrees of freedom; Den = denominator; Sem = effect of semantics.

**Table S37.** Performance indices of the linear-mixed-effects model explaining zRT by semantics.

| AIC | BIC | $R^2$ cond. | $R^2$ (marg.) | ICC | RMSE |
|---|---|---|---|---|---|
| 5356.009 | 5395.511 | 0.891 | **0.011** | 0.889 | 0.371 |

AIC = Akaike information criterion; BIC = Bayesian information criterion; $R^2$ cond. = proportion of variance explained by fixed and random effects**;** $R^2$ marg. = proportion of variance explained by fixed effects (Sem) only; ICC = Intraclass Correlation Coefficient; RMSE = Root Mean Squared Error.

#### Implicit vs. explicit semantic tasks

In our main linear-mixed-model analysis, the semantics variable parsimoniously distinguished semantic and non-semantic experimental conditions. Arguably, semantic conditions can be subdivided into *explicit* semantic conditions (when the task explicitly requires the retrieval of semantic information, e.g. in semantic judgments) and *implicit* semantic conditions (when the task only implicitly probes semantic processing, e.g. in lexical decisions). Therefore, we performed a supplementary linear-mixed-model analysis, where we compared our original “semantics” (semantic vs. non-semantic) model against a “semanticity” (explicit semantic vs. implicit semantic vs. non-semantic) model.

The results show that the “semanticity” model is not better than the “semantics” model (Table S38). In fact, the “semanticity” model is worse (i.e., has a higher AIC) as both models have the same log-likelihood but the “semantics” model is less complex (1 fewer degree of freedom). Therefore, our main “semantics” model remains the optimal model of AG responses across the five studies. Note, however, that it is questionable whether the “semanticity” model is a valid model for our data as only two of the five studies (studies B and C) included both implicit and explicit semantic tasks.

**Table S38**. AIC comparisons to determine the optimal model predicting AG activity.

| Model | ΔLogLik | ΔAIC | df | |
|---|---|---|---|---|
| **PSC ~ Semantics + (1 \| participant) + (1 \| task/condition)** | **5.2** | **0** | **6** | * |
| PSC ~ Semanticity + (1 \| participant) + (1 \| condition) | 5.2 | 2.0 | 7 | |
| PSC ~ 1 + (1 \| participant) + (1 \| task/condition) | 0 | 8.4 | 5 | |

* = winning model; PSC = percent signal change; LogLik: log-likelihood; AIC = Akaike Information Criterion; df = degrees of freedom.
